# BaRTv2: A highly resolved barley reference transcriptome for accurate transcript-specific RNA-seq quantification

**DOI:** 10.1101/2021.09.10.459729

**Authors:** Max Coulter, Juan Carlos Entizne, Wenbin Guo, Micha Bayer, Ronja Wonneberger, Linda Milne, Miriam Schreiber, Allison Haaning, Gary Muehlbauer, Nicola McCallum, John Fuller, Craig Simpson, Nils Stein, John W. S. Brown, Robbie Waugh, Runxuan Zhang

## Abstract

Accurate characterization of splice junctions as well as transcription start and end sites in reference transcriptomes allows precise quantification of transcripts from RNA-seq data and enable detailed investigations of transcriptional and post-transcriptional regulation. Using novel computational methods and a combination of PacBio Iso-seq and Illumina short read sequences from 20 diverse tissues and conditions, we generated a comprehensive and highly resolved barley reference transcript dataset (RTD) from the European 2-row spring barley cultivar Barke (BaRTv2.18). Stringent and thorough filtering was carried out to maintain the quality and accuracy of the splice junctions and transcript start and end sites. BaRTv2.18 shows increased transcript diversity and completeness compared to an earlier version, BaRTv1.0. The accuracy of transcript level quantification, splice junctions and transcript start and end sites has been validated extensively using parallel technologies and analysis, including high resolution RT PCR and 5’ RACE. BaRTv2.18 contains 39,434 genes and 148,260 transcripts, representing the most comprehensive and resolved reference transcriptome in barley to date. It provides an important and high-quality resource for advanced transcriptomic analyses, including both transcriptional and post-transcriptional regulation, with exceptional resolution and precision.

## INTRODUCTION

Barley, the fourth most important cereal crop worldwide, is cultivated across a wide range of environments and is used widely in food and drink for humans and feed for animals (1). Abundant genetic diversity reflects differences in gene expression, whereby developmental, environmental, biotic and abiotic stresses lead to changes in the transcriptome, promoting downstream protein and regulatory responses from the plant (2–5). Such changes occur at the individual transcript level as well as at the gene level, with alternative promotor usage, alternative splicing (AS) and alternative polyadenylation leading to an increased transcript repertoire (6–9).

Detecting changes in gene expression has become routine in plant research using RNA-seq, while identifying transcript variants that are the product of AS is a more recent development (10, 11). A variety of methods exist for RNA-seq data analysis but all require as part of the analysis pipeline either the assembly of transcripts from the dataset (12–14) or an existing reference transcriptome for gene/transcript quantification. Quantification of transcripts with a reference transcript dataset (RTD) can be achieved with speed and precision using established pseudo-alignment programs such as Kallisto and Salmon (15, 16). However, the precision of transcript expression quantification is only as good as the RTD itself, and such resources are currently not available for most plant species (6). The impact of transcript references on quantification accuracy is reflected not only at the transcript-level but also at the gene-level (17, 18). Thus comprehensive, high quality RTDs are key resources for quantitative analysis of gene expression.

Considerable progress has been made in developing plant RTDs using short-read data; first in Arabidopsis (19, 20), and then barley (21). The approaches adopted for their construction were largely based around the accurate detection of splice junctions (SJs) along with the implementation of assembly pipelines that incorporate stringent quality control filters to remove redundant, poorly assembled or fragmentary transcripts (20). During the construction of Arabidopsis AtRTD2, variation in 5’ and 3’ UTR lengths among different transcripts from the same gene were found to skew subsequent transcript quantifications (20). A method for extending transcript ends to the longest transcript in its gene (“padding”) resulted in an overall improvement in quantification accuracy (20) and so was also applied in the creation of an initial barley reference transcriptome BaRTv1.0-QUASI (21). However, close inspection revealed that despite these improvements, incorrect transcripts are still present, and these can be generally attributed to the fact that short read data does not accurately capture transcript 5’ and 3’ ends or phasing of different AS events (6, 22) .

Single molecule sequencing on Pacbio Iso-seq or Oxford Nanopore platforms should in principle overcome issues with 5’ and 3’ end determination and phasing of AS events. Iso-seq has been used to sequence cDNAs of up to 10kb and has been applied to a number of crop species including maize (23), sorghum (24), cotton (8, 25), rubber (26) and tea (27) crops. However, depending on the number of sequencing cycles, some Iso-seq reads suffer from a high sequencing error rate. Other issues include incomplete gene and transcript coverage and high cost per base (28), particularly for data generated from the Sequel platform. Sequencing errors have been addressed by self-correction (29) or hybrid-correction methods (30–32). While these reduce overall error, incorrect clustering of transcripts during self-correction and mis-mapping during hybrid correction can cause over-correction resulting in loss of real transcripts with small AS events, incorrect splice junction determination and generation of new, false splice junctions and transcripts (33). Major challenges remain in accurately determining splice junctions as well as the start and end sites of transcripts.

Defining the 5’ and 3’ ends of transcripts is important in order to identify alternative promotor usage and alternative poly(A) sites. Both of these features are prevalent across biology and add another layer of complexity to regulation of gene expression (8, 34, 35). In Arabidopsis, alternative transcription start sites (TSS) lead to changes in the subcellular localisation of proteins (36) while alternative poly(A) selection has a role in the regulation of processes such as flowering (37). Cap analysis gene expression sequencing (CAGE-Seq) has been used to identify TSS in both Arabidopsis and cotton (8, 34) while polyA-Seq has been used to identify widespread alternative poly(A) sites in the latter (8). Nanopore sequencing has also been used to identify alternative poly(A) sites in Arabidopsis (38–40).

Here we present a new barley reference transcriptome based on the European 2-row spring barley cultivar Barke, making use of its new and accurate TRITEX assembled genome as a reference (41). The new transcriptome combines filtered full-length transcripts from PacBio Iso-seq with stringently filtered Illumina RNA-seq based transcripts. Novel computational methods were used to determine accurate SJs, TSS and TES, which were utilised to generate an Iso-seq-only dataset (BaRT2.0-Iso). In parallel, transcripts from Illumina sequencing of the same barley samples were assembled and stringently filtered to produce a short-read transcript assembly (BaRT2.0-Illumina). Finally, the Iso-seq dataset was supplemented by high quality transcripts from the short-read assembly to overcome the incomplete coverage of some genes in the Iso-seq dataset to generate a new barley reference transcript dataset (BaRTv2.18). Currently BaRTv2.18 comprises 39,434 genes and 148,260 transcripts. High resolution (HR) RT-PCR and a variety of benchmarks demonstrated a significant improvement in BaRTv2.18 compared to BaRTv1.0. Finally, a BaRTv2.18 transcript annotation is provided that includes coding regions and protein variants as well as unproductive transcripts together with functional annotations.

## MATERIALS & METHODS

### Plant material

Twenty one tissues were sampled, all from barley cv. Barke, a European 2-row spring variety with a newly published genome (41). Where possible, tissues were sampled from three or more plants. Third stem internode was sampled from one plant due to the large size of the tissue. Plant material was grown under controlled conditions (16 hours light, 18°C, light intensity 400 µmol.m².s² 8 hours dark, 14°C, 0 µmol.m².s² Humidity 60%) at the James Hutton Institute, or at IPK, Gatersleben. All tissues were flash frozen in liquid nitrogen and added to liquid nitrogen cooled tubes, before being stored at -80°C. Plant material is summarised in Supplementary Table 1 and details of the sample preparations for each tissue can be found in supplementary materials.

### RNA extraction, library preparation and sequencing

RNA extraction was carried out using Qiagen RNeasy Plant mini kits using the manufacturer’s instructions. Total RNA was quality checked using a Bioanalyzer 2100 (Agilent) and RNA samples with the best RIN were used for Iso-seq and RNA-seq library preparation. RNA for Iso-seq and RNA-seq was aliquoted from the same sample. For the seven tissues collected at IPK Gatersleben, RNA was extracted using a Trizol extraction protocol (42) and purified using Qiagen RNAeasy miniprep columns following the manufacturer’s instructions. RNA quality was checked on Agilent RNA HS screen tape.

Single molecule real time sequencing (SMRT) was performed at IPK Gatersleben using the Pacbio Iso-seq method. Full-length cDNA was generated from total RNA using the TeloPrime Full-Length cDNA Amplification Kit V2 from Lexogen (Vienna, Austria). Two separate cDNA synthesis reactions per sample, each with 2 mg total RNA as input, were performed following the manufacturer’s recommendations for Iso-seq library preparation. The purified double-stranded cDNA from both reactions was pooled and amplified in a qPCR reaction to determine the optimal cycle number for large-scale endpoint PCR. The cDNA was amplified in 16 parallel PCR reactions using 2 µl cDNA per reaction with the optimal cycle number using the TeloPrime PCR Add-On Kit V2 (Lexogen). The PCR reactions were purified with AMPure PB Beads according to the post-PCR purification step described in the Iso-seq template preparation protocol for Sequel Systems for libraries without size selection (Pacific Biosciences, Menlo Park, CA). Six of the PCR reactions were pooled to make up fraction 1 enriched in shorter fragments and the other ten PCR reactions were pooled to make up fraction 2 enriched in longer fragments. Equimolar quantities of the two cleaned-up fractions were then pooled and 5 µg of cDNA were used for SMRTbell library preparation using the Sequel 3.0 chemistry (Pacific Biosciences) according to the manufacturer’s protocol. Each library was sequenced on a separate lane of a SMRT cell 1M LR on a PacBio Sequel machine with movie lengths of 1200 min. For tissues collected from IPK, pooled RNA from seven tissues (above) was used to prepare two libraries using the cDNA prepared from TeloPrime v1.0 kit (Lexogen, Vienna, Austria) following the manufacturer’s instructions (41). Libraries were quantified and sequenced on a Pacbio Sequel at IPK Gatersleben (41).

Illumina RNA-seq library preparation and RNA-seq was carried out by Novogene (HK) Company Limited, Hong Kong. The 20 libraries were prepared using NEBNext® Ultra™ Directional RNA Library Prep Kit and sequenced using Illumina NovaSeq 6000 S4 (PE 150). Accession numbers for accessing the raw data generated for each tissue are provided under Data Availability.

### Iso-seq processing

#### Generation of the initial transcript assembly

The 20 Iso-seq libraries were processed individually. Raw subreads were initially processed using the CCS tool from IsoSeq3 to create circular consensus sequences (CCS). The resulting CCS reads were stripped of adapter sequences and poly-A tails using lima and refine from Isoseq3. A further clean-up of PolyA tails was carried out using the Transcriptome Annotation by Modular Algorithms (TAMA) tama_flnc_polya_cleanup.py script. The resulting reads were mapped to the Barke genome (41) using Minimap2. TAMA collapse (https://github.com/GenomeRIK/tama/wiki/Tama-Collapse) was used to process the read mappings for each sample to generate a non-redundant set of transcripts. The TAMA collapse output files from 20 samples were merged together using TAMA merge (https://github.com/GenomeRIK/tama/wiki/Tama-Merge). Details of the parameters used for each step can be found in the Suppl. materials.

#### Identification of high confidence splice junctions

Sequence errors in Iso-seq reads can lead to the identification of incorrect SJ co-ordinates. A collection of high confidence SJs was selected where they had 1) canonical donor and acceptor sequence motifs and 2) had at least one supporting read that had a perfect match within 10 nucleotides upstream and downstream of the SJ (43). Splice junction and read information was obtained from the TAMA collapse output using a customised script (https://github.com/maxecoulter/BaRT-2) for each of the 20 libraries. In addition, a dataset of high confidence splice junctions was created from Illumina short read RNA-seq data using the output from the mapping tool STAR. High confidence SJs were selected based on having 1) canonical motifs; 2) an overhang of >=10 nt and 3) more than 5 supporting reads.

To remove splice junctions that were potentially caused by template switching (44), the Hamming distance was calculated between the last 8 bases of the splice junction’s upstream exon and the last 8 bases of the intron for each splice junction. The same was done for the first 8 bases of the intron and the first 8 bases of the downstream exon for each splice junction. A Hamming distance of 1 or less (1 nucleotide difference between the two 8 nucleotide sequences being compared) in either comparison was considered a potential template switching event, and these splice junctions were removed from high confidence SJs.

#### Identification of Transcript Start Sites (TSS) / Transcript End Sites (TES)

For 3’ ends, reads potentially with internal priming (45) were identified from the poly(A) output from TAMA collapse. Reads/transcripts with >16 As in the 20 nucleotides following the 3’ end in the internal gene sequence in the genome were considered potential off-priming and were removed. For many genes, there were long reads representing full-length transcripts but also 5’ and 3’ truncated reads likely due to degradation or incomplete cDNA synthesis. To distinguish between real and false transcript end sites, we used two methods depending on expression level (43). We assumed that for genes with multiple reads with different ends, the start and ends of fragmentary reads would be distributed randomly and uniformly between the 5’ and 3’ ends, while true TSS/TES would have more read support than what would be expected by random. For each gene, we used a binomial discrete mass function to infer the probability that the read start and end position was likely to be random using the following formula (46) :

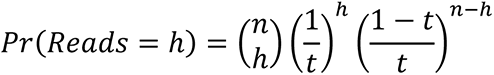

where *n* is the total number of reads in a gene, *ℎ* is the number of reads with a start or end site being tested and *t* is the total number of starting or end sites in the gene. TSS or TES within a gene were considered to be high confident (HC) for downstream processing when the probability of having *ℎ* reads at the site by random < 0.01.

To increase the gene coverage for low abundance genes (typically < 10 reads) without at least one significantly enriched TSS and TES, we retained TSS/TES if they had start and end support from 2 or more reads within a sliding window of +/-20 and +/-60 for starts and ends respectively.

#### Filtering the poorly supported transcript assembly

The transcripts from the initial assembly were filtered using the datasets of HC SJ and HC TSS/TES sites. For genes with HC TSS and TES, transcripts were kept if a) their starts locate within 10 nucleotides of the defined HC TSS and their ends locate within 30 nucleotides of the defined HC TES and b) all splice junctions in the transcript were present in the list of HC Iso-seq splice junctions. To remove redundancy due to variation at the 5’ and 3’ sites, transcripts were run through TAMA merge to create BaRTv2.0-Iso, using the settings “m 0 -a 100 -z 100”.

#### Illumina RNA-seq processing and analysis

In conjunction with FastQC (http://www.bioinformatics.babraham.ac.uk/projects/fastqc/) Trimmomatic version 0.39 was applied to the raw Barke RNA-seq reads (47) for quality control and adapter trimming. The trimmed RNA-seq reads were aligned to the Barke reference genome with STAR (Spliced Transcripts Alignment to Reference) (48). A two-pass method in which the splice junction outputs of a first STAR pass were used for indexing a second pass, was applied to improve the accuracy and sensitivity of read alignment and splice junction identification. Minimum and maximum intron sizes were 60 and 15,000 for the alignment. Mismatches were not allowed. Transcripts were assembled from the read alignments with three tools: Cufflinks version 2.2.1 (49), StringTie version 2.0 (13) and Scallop version 0.10.4 (50) to take advantage of complementarity among the tools in capturing the diversity of transcripts. One transcript dataset was generated for each sample by each assembler. The sixty individual assemblies were merged and quality filtered into a single transcript assembly (BaRTv2.0-Illumina) with RTDMaker (https://github.com/anonconda/RTDmaker). RTDmaker identifies and removes 1) redundant transcripts; 2) transcripts with splice junctions with low read support; 3) transcript fragments; 4) poorly supported antisense transcripts; 5) unstranded models; 6) antisense transcript fragments and 7) low expressed transcripts (TPM cut-off default = 1).

#### Merging BaRTv2.0-Iso and BaRTv2.0-Illumina

Transcripts from the high confidence Iso-Seq dataset (“BaRTv2.0-Iso”) and Illumina dataset (“BaRTv2.0-Illumina”) were initially merged with TAMA merge using “wobble” parameters of -m 0 -a 100 -z 100 (51). This allowed the removal of redundant transcripts with exact matches for splice junctions with small variable ends (100 bp). During the merge, priority was given to 5’ and 3’ ends of Iso-Seq transcripts by setting “capped” flag for Iso-Seq dataset and “no_cap” for BaRTv2.0-Illumina dataset.

Custom code was used to carry out further filtering according to the following principles: a) BaRTv2.0-Iso was considered to be the most accurate transcriptome assembly, so all transcripts with HC TSS/TES were kept; b) Iso-seq transcripts without HC TSS and TES were only kept if they had Illumina support and were at least 80% of the length of the average length of Illumina transcripts in that gene; and c), Illumina transcripts were retained for genes with no Iso-seq coverage or if they contained novel SJs.

Finally, to remove the high number of mono-exonic genes likely due to DNA contamination (51), the mono-exonic genes were re-analysed using expression abundance and those genes with expression higher than 1 TPM in less than 2 samples were removed (over 13.4k genes). To keep bona fide single exon protein-coding genes that were tissue-, organ- or condition-specific, 2.4k single exon protein-coding genes with ORFs greater than 100 amino acids predicted by Transuite (52) were kept.

#### Motif enrichment analysis

To explore motifs commonly associated with TSS and TES (Supplementary Table 2), for every predicted transcript 5’ and 3’ site in BaRTv1.0 and BaRTv2.18, the sequence ±550 nucleotides either side of the TSS or TES was extracted from the genome, and after a regex search the position coordinates of all matching motifs relative to the sequence were extracted. As a control, the same number of random sites were taken from random chromosomes, and the above analysis was carried out.

#### High resolution RT PCR

RNA from12 of the tissues used to construct BaRTv2.18 was used for high resolution (HR) RT-PCR validation. A list of samples is provided in Suppl. Table 4. 47 primer pairs used for covering 99 RT-PCR products designed previously for BaRTv1.0 (21) were identified where 1) the primer pair had perfect matches to the Barke genome and 2) the interval between the primer pairs had good RNA-seq coverage (>1,000 reads in at least two samples) identified with samtools bedcov (53). The full list of primers, products and RT-PCR product proportions used in the analysis are in Supplementary Table 3A, B and C. 5’ labelling, first strand cDNA synthesis, HR RT-PCR and detection were carried out as described previously (21, 54). To compare with BaRTv1.0, taking into account possible differences in sequence, expression and AS due to genotypic variation, 86 primer sets (Suppl. Tables 5 and 6) from the HR RT-PCR dataset of Flores et al., 2019 (21) were also used, and the data reanalysed according to the method described in (21). A detailed description of the method along with the custom code used to carry out the analysis is described at https://github.com/maxecoulter/BaRT-2, and in the supplementary methods section.

#### 5’RACE

Transcription start sites were characterised by cloning and sequencing 5’ RACE products using the SMARTer RACE 5’/3’ kit (Takara Bio USA, Inc). In summary, antisense gene-specific primers were designed downstream of predicted Iso-seq TSS (Supplementary Table 4). RNA samples from Barke inflorescence and peduncle were pooled and 1µg was used in the 5’ RACE assays. After reverse transcription, 3’-end tailing and annealing of the SMARTer II A oligonucleotide, the second strand was synthesised to form a double stranded cDNA. RACE products were amplified using the universal primers and the gene-specific primers (Supplementary Table 4), followed by in-fusion cloning into the vector pRACE using the 15 base overlap designed into the gene-specific primer. DNA from selected plasmids was extracted using Wizard Plus SV minipreps DNA purification system (Promega) and sequenced using the M13F primer. Sequences were aligned to the relevant gene transcripts in the BaRTv2.18 release using Clustal Omega on the EMBL-EBI platform (55).

#### Annotation

Transuite 0.1.2 was used to identify ORFs in BaRTv2.18 transcripts (52) and the predicted protein sequences used for annotation using PANNZER2 (56) to produce predicted functions and GO terms. A minimum Positive Predictive Value (PPV) of 0.3 was used to include a predicted annotation in the dataset. TranSuite is a program which identifies coding and non-coding transcripts, generates accurate translations of transcripts and identifies features of AS and Nonsense mediated mRNA decay (NMD). SUPPA2 version 2.3 was used to characterise alternative splicing events that occur within BaRTv2.18 (57).

#### Benchmarking

Transcript lengths were computed from the candidate FASTA files using samtools faidx v. 1.9 (53). The number of duplicated sequences was established using seqkit rmdup v0.12.0 (58). The number of genomic gaps denoted by Ns in the candidate transcripts was counted using custom Java code. BUSCO v4.0.6 (59) was used to benchmark transcript set completeness and transcript fragmentation.

To establish the percentage of assembled transcripts that were chimeric (either as a result of readthrough or bioinformatics artefacts), GMAP version 2018-07-04 (60) was used to map the Haruna Nijo full length cDNA (flcDNA) sequences (61) to the reference genomes of cultivars Morex (62) and Barke (41), using the parameters “-- min-identity=0.96” and “--min-trimmed-coverage=0.95”. The resulting GFF output was converted to GTF format using gffread v0.11.6. (http://ccb.jhu.edu/software/stringtie/gff.shtml#gffread). Using both reference genomes was intended to prevent variation in genome composition from affecting the outcome of the mapping, as the previous version of BaRT was based on the Morex genome. The resulting GTF files were then screened for positional overlap with the GTF files of the candidate assemblies using bedtools intersect (63), using the flcDNA mapping to the reference genome underlying the candidate assembly (BaRTv1.0 = Morex, BaRTv2.18 = Barke). Custom Java code was used to parse the output from bedtools intersect and count the number of flcDNA sequences spanning a candidate transcript, with those candidate transcripts overlapping more than one flcDNA being counted as chimeric.

## RESULTS

### Construction of the BaRTv2.0 Iso-seq transcriptome (BaRT2.0-Iso)

To maximise the number and diversity of transcripts, 21 samples from a range of tissues, growth stages and treatments were used (Supplementary Table 1). The workflow for the Iso-seq data processing is described in Figure 1. Briefly, Iso-seq reads were pre-processed to full-length non-chimaeric reads (FLNC) and mapped to the Barke genome before being filtered against high confidence SJ, TSS and TES datasets to generate the Barke Iso-seq transcriptome, BaRTv2.0-Iso (Figure 1A). This latter step is described in more detail in Figure 2. In parallel, Illumina reads were generated from the 20 samples, pre-processed and transcripts assembled with three different assemblers (Figure 1B). Assemblies were merged and processed to give the short read assembled transcriptome, BaRTv2.0-Illumina (Figure 1B). Finally, the Iso-seq and Illumina datasets were merged and filtered to remove redundancy and prioritise Iso-seq transcripts over Illumina transcripts. The resultant RTD, BaRTv2.10 underwent a comprehensive series of final filtering steps to, for example, remove low expressed mono-exonic transcripts to give the final barley (Barke) RTD, BaRTv2.18. The production of BaRTv2.18, generation of HC SJ, TSS and TES datasets and the quality control steps are described below.

**Figure 1:**
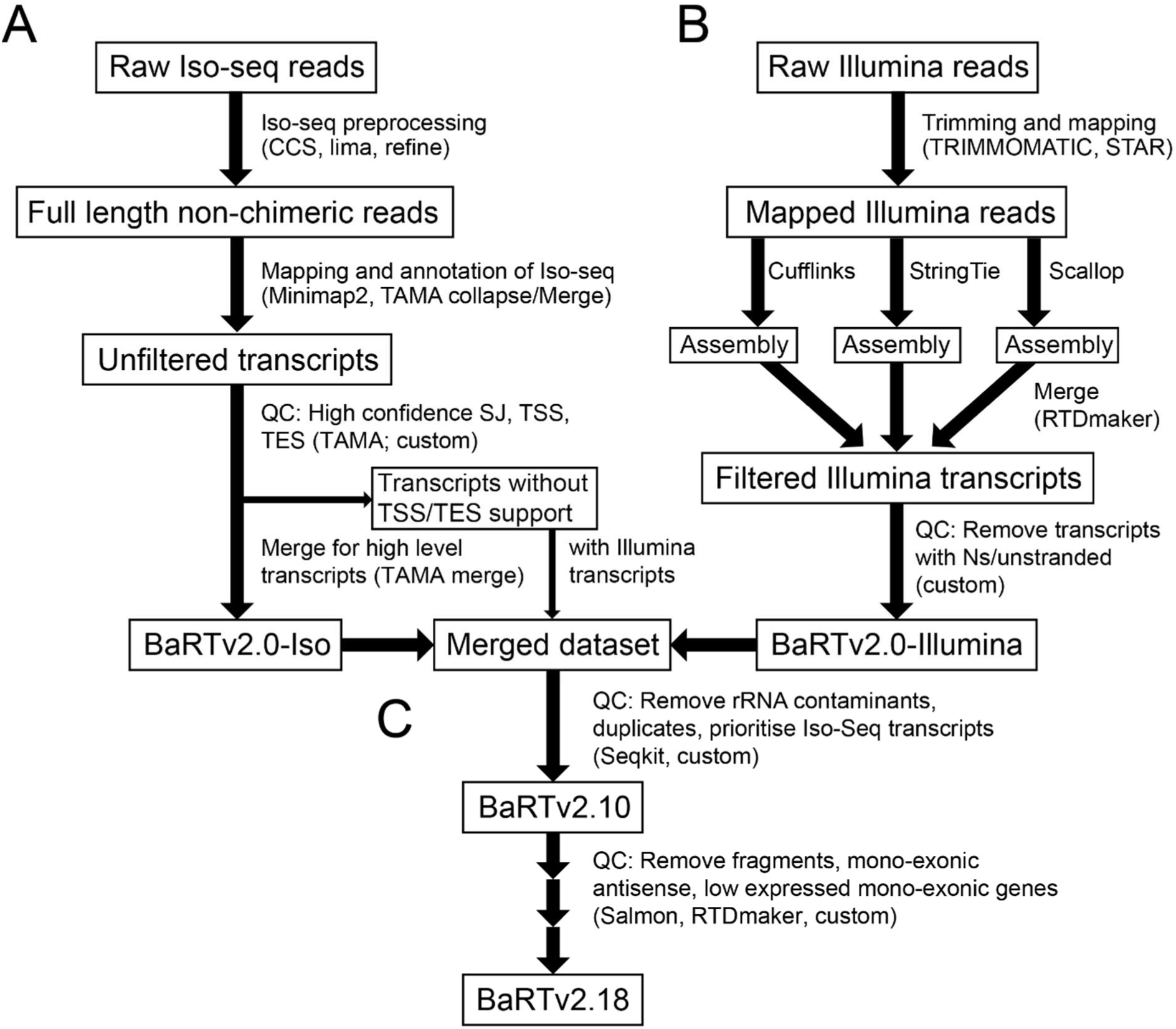
Outline of the pipeline used to generate BaRTv2.18. Iso-seq and Illumina datasets were processed separately (**A, B**) and then combined at the final stage (**C**). Software and scripts are indicated in brackets, with “custom” referring to custom code. **A**) For the Iso-seq dataset, raw subreads were pre-processed using Isoseq3 software (CCS, lima and refine) to create full length non chimeric reads. These reads were mapped to the Barke genome using Minimap2. Mapped reads were used as input for TAMA collapse and TAMA merge to create a set of unfiltered transcripts. Filtering was carried out using custom codes and code described in Figure 2, to remove fragments, unsupported splice junctions and TSS/TES. Transcripts from genes with low read end support that had HC splice junctions were kept in a separate file for potential inclusion depending on similarity to Illumina transcripts. Redundant transcripts were removed using TAMA merge to create BaRTv2.0-Iso. **B**) Illumina reads were trimmed using Trimmomatic and mapped to the Barke genome using STAR. Transcripts were assembled using three separate assemblers: Cufflinks, Stringtie and Scallop. The resulting assemblies were merged and filtered using the RTDmaker software. The resulting annotation was further filtered to remove transcripts overlapping Ns and transcripts with no strand information to create the BaRTv2.0-Illumina dataset. **C**) The BaRTv2.0-Iso and BaRTv2.0-Illumina datasets were merged using TAMA merge (giving priority to BaRTv2.0-Iso based transcripts). Further filtering was carried out using custom code to remove redundant Illumina transcripts. Duplicate transcripts were removed using seqkit rmdup, as well as potential rRNA. The resulting transcriptome BaRTv2.10 went through further filtering steps with RTDmaker including removal of low expressed mono-exon genes based on Salmon quantifications to generate the final transcriptome BaRTv2.18.

**Figure 2:**
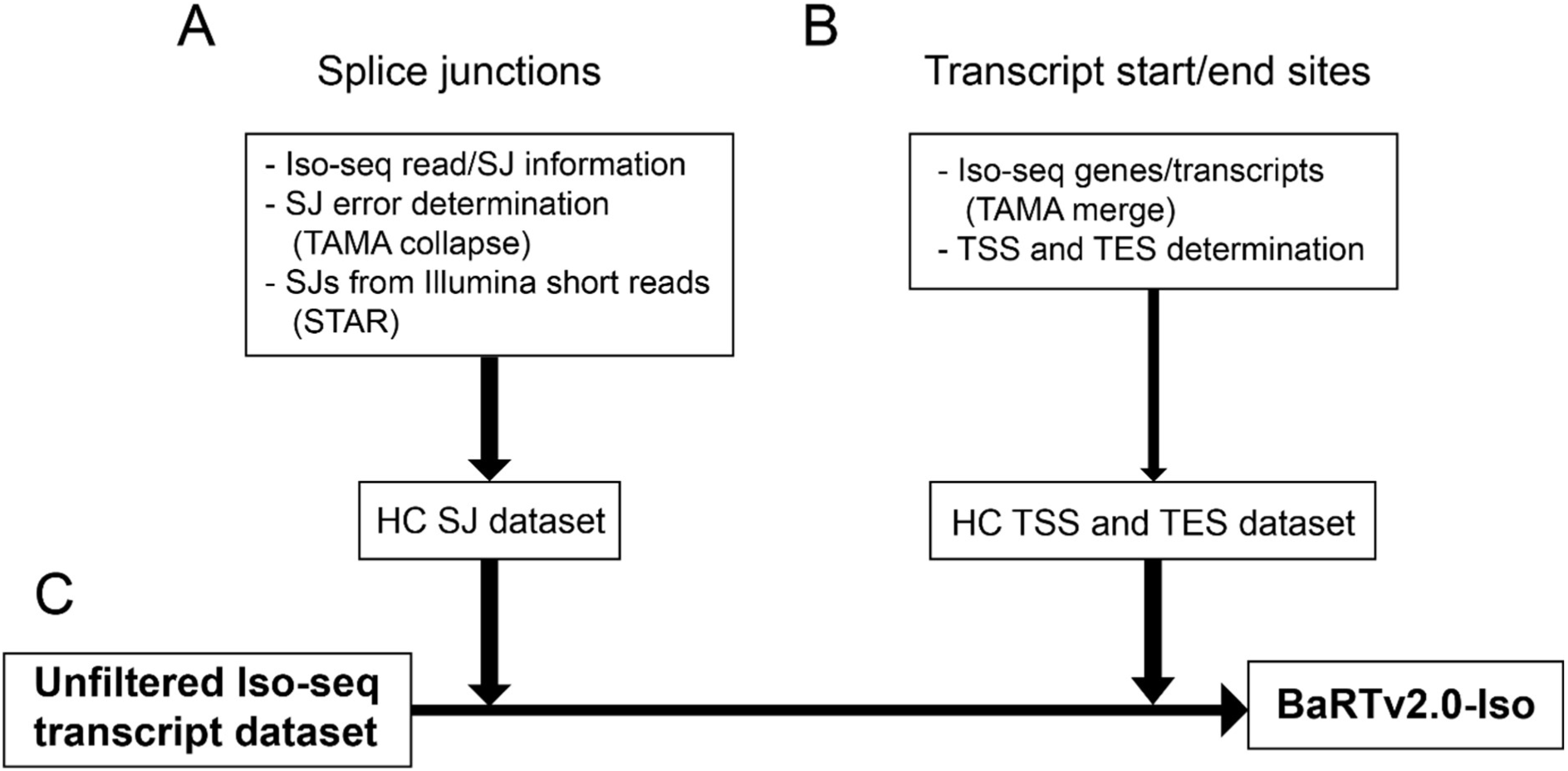
Overview of generation of BaRTv2.0-Iso using HC SJ, TSS and TES datasets. **A**) The high confidence dataset of SJs was generated from Iso-seq data using the error distribution profiles around SJs to remove false SJs and from Illumina SJ data. **B**) The HC TSS and TES dataset was generated from Iso-seq data using two methods dependent on transcript abundance. **C**) The unfiltered Iso-seq transcript dataset was filtered against the HC SJ, TSS and TES datasets and only transcripts with HC information were retained in BaRTv2.0-Iso.

For Iso-seq, the first stage processed raw Iso-seq reads with the PacBio Iso-seq 3 pipeline to yield a combined total of 8,113,088 CCS reads with a mean of 405,654 reads per sample. After demultiplexing, removal of polyA tails and concatemers, a total of 7,395,557 FLNC reads were generated of which 93.7% (6,930,934) mapped to the Barke genome with a mean of 346,546 mapped FLNC reads per sample (Figure 1A; Supplementary Table 5). For the second stage of the analysis we used the TAMA suite of programs (33). “TAMA collapse” integrated redundant transcript models in each sample providing 20 initial annotation files. These were then merged with TAMA merge (merges multiple transcriptomes while maintaining source information) to create an unfiltered transcripts dataset containing a total of 33,550 genes and 2,004,544 transcripts with an average of 15,733 genes and 166,048 transcripts per sample (Figure 1A; Supplementary Table 5). At this stage transcripts were defined at 1bp resolution (I.e. any small differences in mapping between two reads would lead to two transcripts being annotated).

### Determination of high confidence Splice junctions (SJ), Transcription Start Sites (TSS) and Transcription End Sites (TES)

Major challenges remain in the accurate definition of splice junctions (SJs) and transcript start and end sites (TSS and TES) due to sequencing errors and degradation of mRNAs in vivo or during processing. We applied newly developed methods of Iso-seq analysis which use sets of high confidence SJs and TSS/TES to define these key features (43, 51). Some FLNC reads still have relatively high error rates (ca. 3% - (33)). Mapping FLNCs to the genome can be inaccurate due to the phenomenon of edge wander (64) where inaccuracies in defining the boundaries of a gap are the main source of error, for example when aligning a sequence with spliced intron to a genomic reference. Mis-mapping of reads containing spliced introns is exacerbated by local sequence errors (± 10 nt) that frequently generate false SJs leading to transcripts with incorrect intron-exon boundaries (43). To address this issue, a high confidence (HC) SJ dataset was created based on both Iso-seq and short read RNA-seq data (see below). A total of 257,496 SJs were identified in the Iso-seq dataset of which 164,860 were designated HC (64%) (Supplementary Table 6A). Of the 92,636 SJs which were filtered out, 31,541 contained mis-matches within 10 bp of a splice junction, 2,687 were template switching events and the rest (58, 408) had non-canonical splice site dinucleotides (Supplementary Table 6B). Template switching events are an artefact of cDNA synthesis which can lead to exonic sequence loss and incorrect SJ designation (44). Iso-seq transcripts that only contains SJs in the HC dataset are kept for downstream processing (see below; Figure 1A; Figure 2A).

Despite applying the Teloprime cap capture system to enrich for full length Iso-seq reads, numerous 5’ and 3’ truncated reads were observed in our dataset that likely reflect degradation of transcripts either in the cell or during RNA preparation. As for SJs, datasets of HC TSS and TES sites were generated from the Iso-seq data (Figure 2B). An important consideration is that the read abundance of genes in the Iso-seq data covered a large dynamic range with, for example, over 8,000 genes having only one or two FLNC reads while the top 10% of expressed genes contained 79% of FLNC reads in the dataset. The identification of HC TSS/TES sites for high and low expressed genes presents very different issues. For example, for highly expressed genes, the major challenge is to reduce false end sites from relatively high numbers of transcript fragments (degradation products) while identifying the dominant, bona fide sites. For low abundance genes, the problem is to obtain enough experimental evidence to define a TSS/TES.

We used two methods to determine TSS and TES depending on the level of expression. Firstly a binomial probability (46) was used on all genes in the dataset to identify TSS and TES sites that were significantly enriched in read numbers relative to the total number of reads from the gene (Figure 3A). That is, bona fide TSS and TES will occur more frequently and will be significantly enriched whereas the randomly distributed ends derived from degraded mRNAs are unlikely to be significantly enriched and can be removed. We found that many transcripts when compared had small variations in their 5’ or 3’ ends; such variation is likely to reflect the stochastic nature of most TSS/TES that show a distribution around a dominant site (34). To account for this during determination of TSS and TES, we examined the distribution of read end sites around all end sites with high (>10 reads) support. The majority of transcript 5’ ends occurred within a window ±10 bp of the major site while 3’ ends were more variable with the majority of them within a window of ±30 bp (Supplementary Figure 1A, B). These windows were applied to allow for stochastic variation in transcript ends while removing transcripts with false ends (Figure 3A). A total of 43,302 and 59,944 significantly enriched TSS and TES, respectively were identified in 14,589 genes (Supplementary Table 7A).

**Figure 3:**
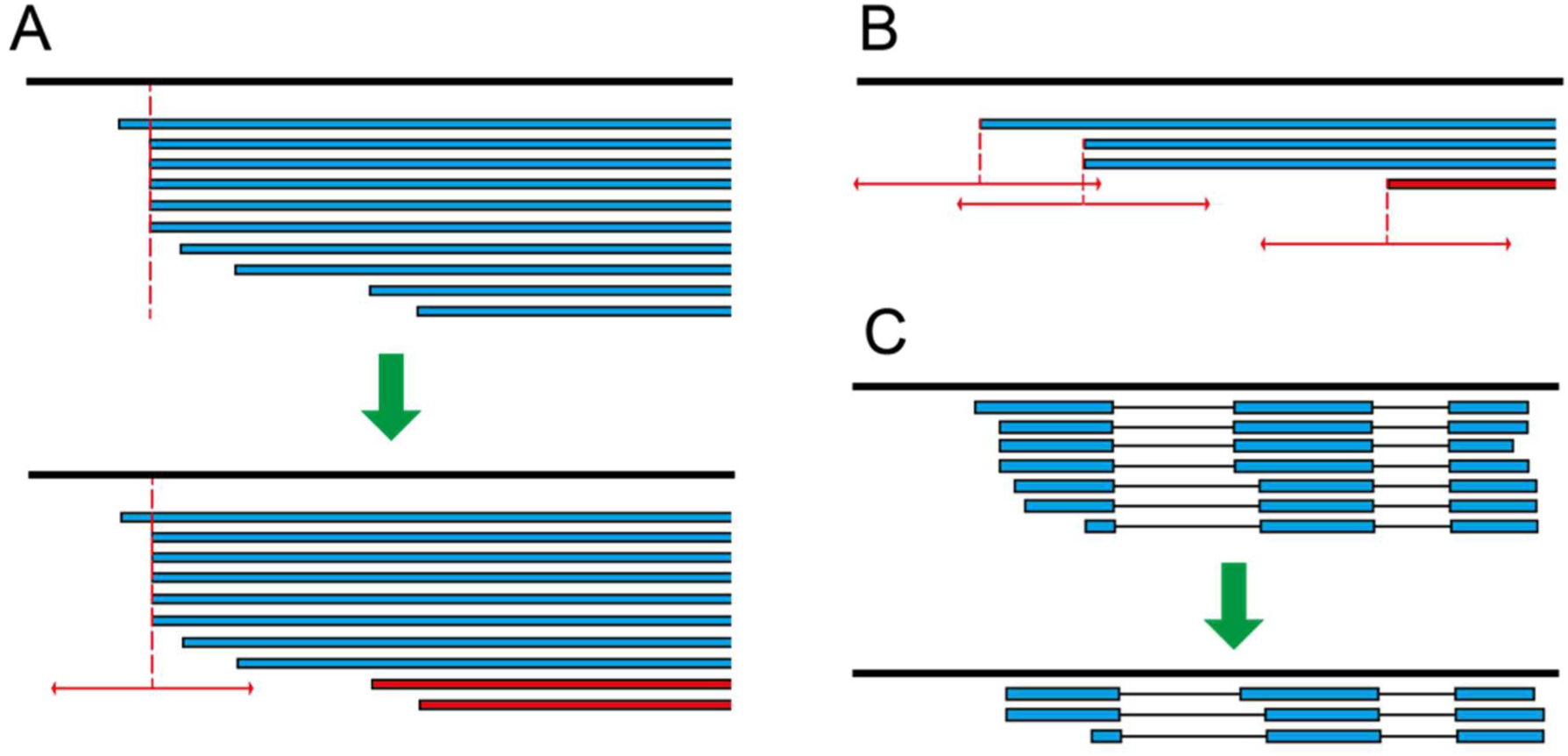
Schematic diagram of filtering and merging methods used to generate BaRTv2.0-Iso. Transcripts or transcript ends (blue) are shown aligned to the genome (black line). **A**) Binomial enrichment method: transcripts with significantly enriched end sites (top panel, dotted line) and other transcripts with similar end sites within a window (+/-10bp for TSS, +/-30bp for TES sites) (horizontal arrows) were kept (**A**, bottom panel). Transcripts with ends outside of the window from an enriched site were removed (red). **B**) For genes with lower Iso-seq coverage, the fixed window method was used where transcripts whose ends fell within a sliding window (+/-20bp for TSS, +/-60bp for TES) (horizontal arrows) were retained while those with unsupported ends were removed (red). **C**) TAMA merge was used after Iso-seq filtering to merge transcripts with same intron coordinates and similar ends (within +/-100bp) to the length of the transcripts with highest abundance ends.

For genes where the binomial probability approach failed to reveal enriched TSS/TES (normally due to the low expression of that gene), we require at least two reads/transcripts had to have 5’ or 3’ ends within a window of ± 20 bp and ± 60 bp, respectively (Figure 3B). These windows were determined based on the distribution of ends around major end sites shown in Supplementary Figure 1. By applying these criteria, HC TSS/TES were identified for a further 10,041 genes (Supplementary Table 7A). Finally, the remaining 8,739 genes had neither HC TSS nor TES sites predicted by either method. These genes had either a small number of mapped FLNC reads without closely related ends or had only one FLNC read supporting (Supplementary Table 7B). As many of these genes had transcripts containing HC SJs, their transcripts were retained in a separate .bed file for comparison to Illumina short read assembled transcripts to provide additional information on transcript features and, in some cases, were re-integrated into the RTD (Figure 1A; Supplementary Table 7B).

### Generation of Iso-seq transcripts in BaRT2.0-Iso

The datasets of HC SJs, HC TSS and TES were used to quality control the Iso-seq transcript dataset of 2,004,544 transcripts created using TAMA merge at single nucleotide resolution (Figure 1A; Figure 2C). Only transcripts containing HC SJs, TSS and TES were retained giving 1,134,325 transcripts from 24,566 genes. Many of the differences among transcripts from the same gene were minor, with transcripts containing similar (but not identical) TSS and TES. To remove redundancy due to such variation in TSS and TES, transcripts were again run through TAMA merge. The settings “m 0 -a 100 -z 100” ensured that transcripts with the same set of SJs and with similar 5’ and 3’ ends within 100 bp were merged together to the most abundant TSS and TES sites (Figure 3C). The final BaRTv2.0-Iso dataset contained 103,330 transcripts (reduced from 1,134,325) from 24,630 genes (Figure 1A).

### Assembly of the Illumina transcriptome (BaRT2.0-Illumina)

Transcripts were assembled from alignments of the 20 short read libraries (Figure 1B; Supplementary Table 1). Illumina reads Mapped by STAR from each library were assembled with three tools: Cufflinks (49), StringTie (13) and Scallop (50) yielding one transcript assembly per sample per assembler (Figure 1B). The 60 individual assemblies were merged with RTDmaker (https://github.com/anonconda/RTDmaker) to give an Illumina transcript dataset. The amalgamated transcript assembly was quality controlled using RTDmaker which is an automated and highly refined extension of the original protocol implemented to generate Arabidopsis AtRTD and AtRTD2 (19, 20). From the original total of >3.6M transcript models, 3,460,323 were rejected using the above criteria (Supplementary Table 8). The resulting BaRT2.0-Illumina transcriptome consisted of 54,017 genes and 142,174 transcripts.

### Merging Iso-seq and Illumina datasets

Gene, transcript and splice junction identity of the BaRT2.0-Iso and BaRT2.0-Illumina transcriptomes were initially compared using TAMA merge with parameters where SJs had to be exact matches, and TSS/TES located within 100 bp. The overlap between Iso and Illumina genes was high, with 23,058 (95.1%) of the Iso-seq genes also present in Illumina data (Figure 4A). Only 1,182 genes were unique to the Iso-seq dataset while 32,150 genes were unique to the Illumina assembly illustrating the lower gene coverage in the Iso-seq dataset. For SJs, 104,086 were common to both datasets (Figure 4B). The SJs unique to BaRT2.0-Illumina are largely from genes missing in the Iso-seq dataset. The 23,313 SJs unique to BaRT2.0-Iso potentially reflect novel transcript isoforms. In contrast to the SJ and gene overlaps, the transcript overlap was relatively low (Figure 4C). Only 7,024 transcripts shared the same transcript structures between the Illumina and Iso-seq datasets. The transcript differences reflect 5’ and 3’ end variation, AS isoforms unique to one or other platform, transcript fragments and possible mis-assembly of some short read transcripts. Many of these differences were resolved during merging and subsequent quality control. To examine 5’ and 3’ end differences, we identified transcripts with identical SJ sets in both BaRT2.0-Iso and Bart2.0-Illumina. Of the 41,079 transcripts, 34,055 (82.3% had ends differing by >100 bp with only 7,024 (Figure 4C) having ends within the 100 bp window. Thus, as expected, most of the defined TSS and TES in Iso-seq transcripts did not have matches among the short read assembled transcripts.

**Figure 4:**
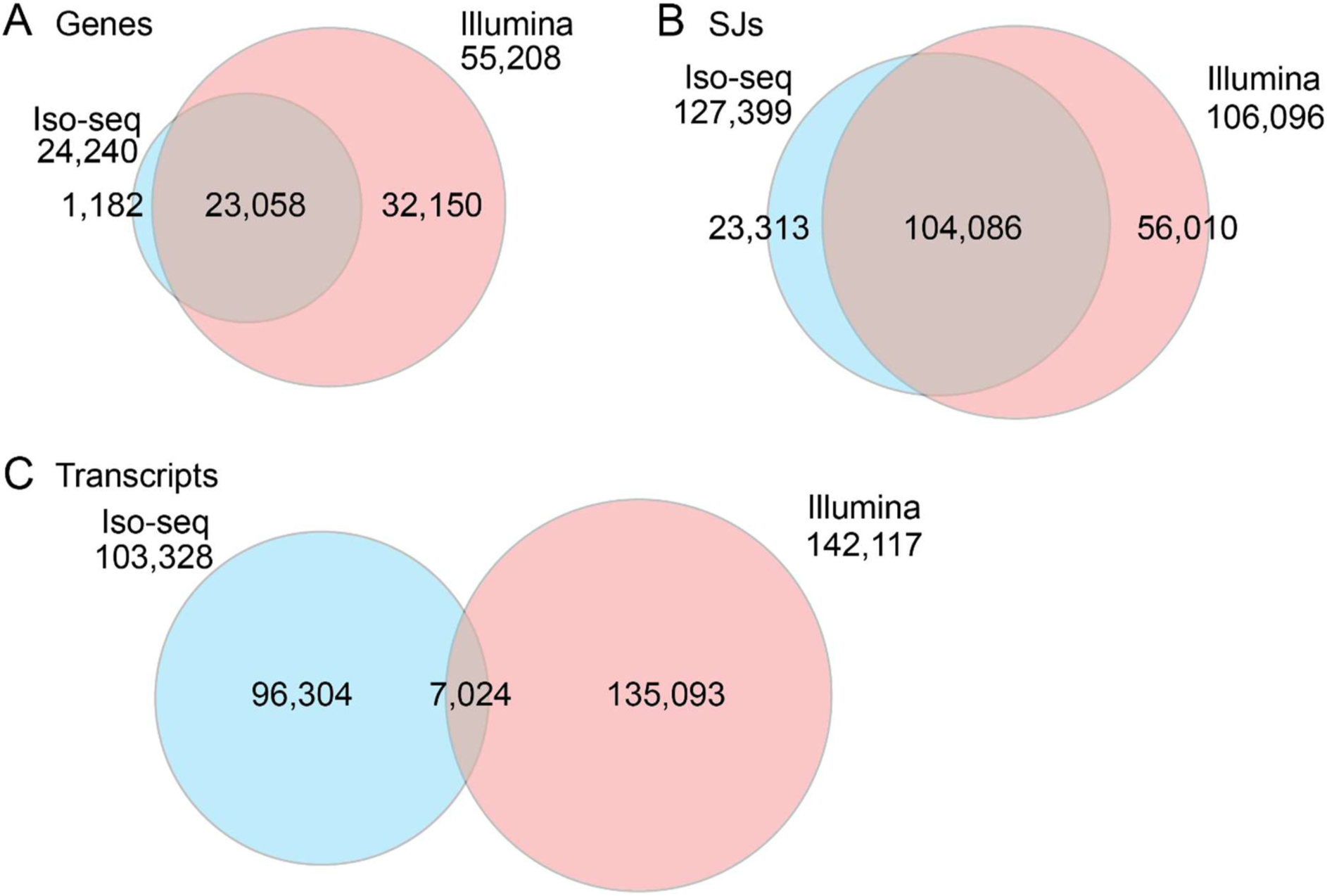
Venn diagrams showing overlap between genes, SJs and transcripts produced by Iso-seq and Illumina. A) Genes, B) splice junctions and C) transcripts in the merged Iso-seq/Illumina dataset before filtering to create BaRTv2.18.

**Figure 5:**
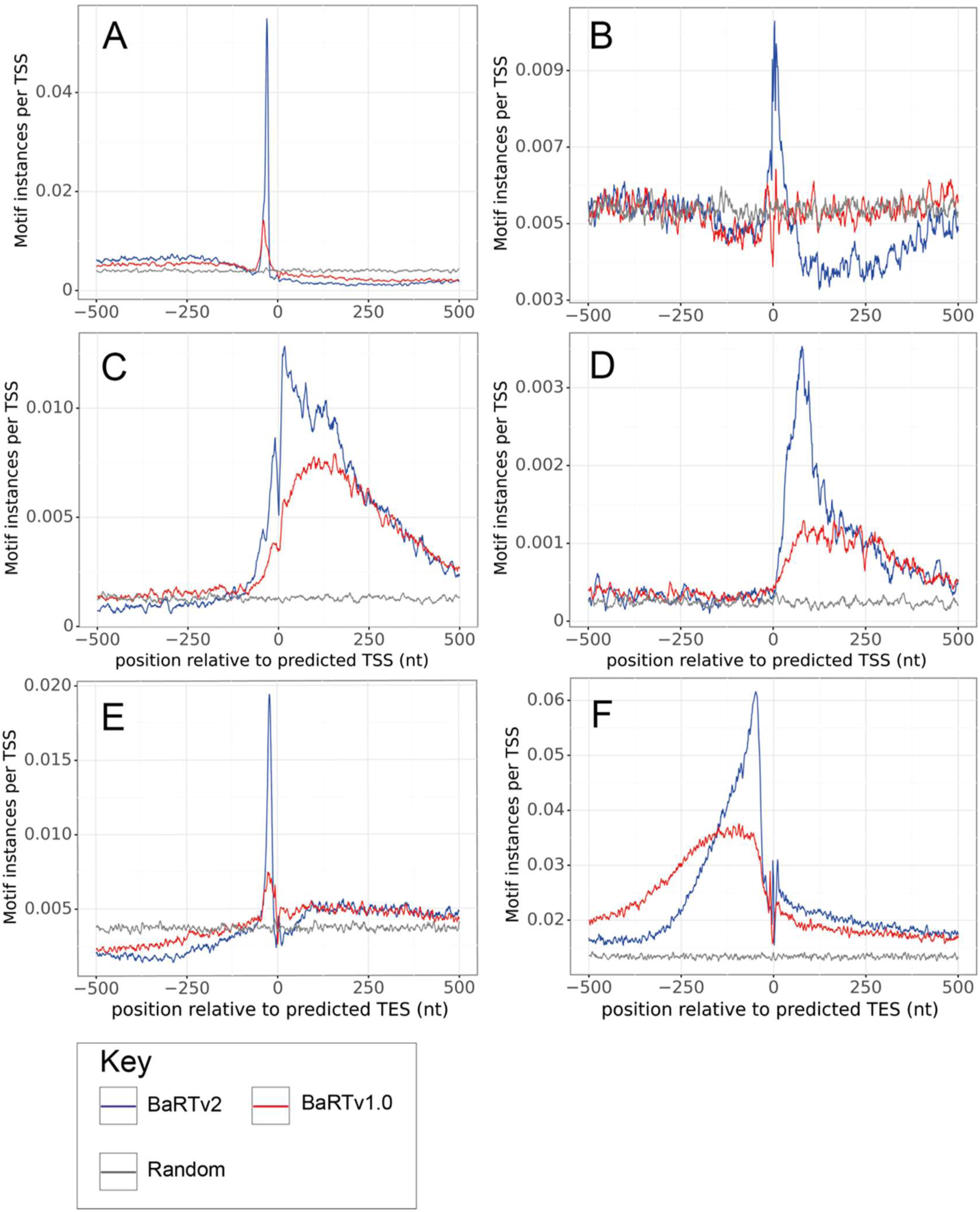
Enrichment of well-known sequence motifs associated with TSS and TES. (**A-D**) TSS-associated motifs and (**E-F**) TES-associated motifs. Lines indicate the frequency of instances of each motif in relation to the end sites with red lines showing results from BaRTv1.0 and blue lines from BaRTv2.18; grey lines represent random control. TSS motifs: **A** TATA box, **B** Inr, **C** Y-patch, **D** Kozak motif; TES motifs: **E** PAS and **F** CFlm.

In order to create a single reference transcript dataset, the BaRTv2.0-Iso and BaRTv2.0-Illumina assemblies were merged and filtered using custom scripts to remove redundant transcripts and a) prioritise Iso-seq transcripts as they contained HC SJs, TSS and TES; and b) retain Illumina transcripts that represent genes absent from the Iso-seq dataset or which contained novel SJs (Figure 1C). This merging and filtering resulted in the development of an initial integrated reference transcript dataset (BaRTv2.10), which contained a total of 56,029 genes and 173,635 transcripts. BaRT2.10 contained an unexpectedly high number of mono-exonic genes including mono-exonic antisense genes, as well as some transcript fragments. The vast majority of the mono-exonic genes were derived from the short-read Illumina assembly while some fragments and redundant transcripts still remained in both the Iso and Illumina assemblies. BaRT2.10 was processed through RTDmaker in a series of steps to remove transcript fragments and mono-exonic genes with low expression. A high number of mono-exonic genes had low expression and were only observed in a single sample, likely due to DNA contamination of samples. Thus mono-exonic genes with expression detected in less than two samples were identified and removed unless they were protein coding genes. The final barley RTD version, BaRTv2.18 consisted of 39,434 genes and 148,260 transcripts (Figure 1C; Supplementary Table 9).

We also compared the frequency of different AS event types in BaRTv2.18 using SUPPA2 (57). The frequency of different AS events is similar to other plant species, with more alternative 3’ splicing events (24.8%) than alternative 5’ splicing events (16.4%) and relatively few exon skipping events (11.7%) (64). Intron retention (IR) is far more frequent in plants than in animals with around 40% of plant AS events being IR (65). Intron retention events made up 41.5% of AS events in BaRTv2.18 (Supplementary Table 10).

### Characterisation of BaRTv2.18 genes and transcripts

The genes and transcripts in BaRTv2.18 were characterised using TranSuite. Transuite outputs detail on protein-coding and non-coding genes and transcripts, protein-coding and unproductive transcripts from protein-coding genes, a breakdown of genes by exon number (single versus multiple) and number of isoforms (single versus multiple), NMD predictions and translations of all transcripts in the RTD. These results are summarised in Figure 6 and Supplementary Tables 9 and 11. Eighty-one percent of genes in BaRTv2.18 coded for proteins and 19% were non-protein-coding genes (Figure 6A; Supplementary Table 9) with about 70% containing more than one exon and 30% single exon genes (Figure 6B; Supplementary Table 9). Of the multi-exonic genes, 73% had more than one transcript isoform and 27% produced a single transcript; the mono-exonic genes had single transcripts (Figure 6C; Supplementary Table 9). For protein-coding genes only, nearly 58% were multi-exonic with more than one transcript isoform and so are AS, agreeing with previous estimates of the prevalence of AS in Arabidopsis and other species (20, 65, 66). The 7,501 non-protein-coding genes generated 11,723 transcripts; 2,160 genes were multi-exonic (spliced) but with a single transcript and 1,179 genes were alternatively spliced (Supplementary Table 9).

**Figure 6.**
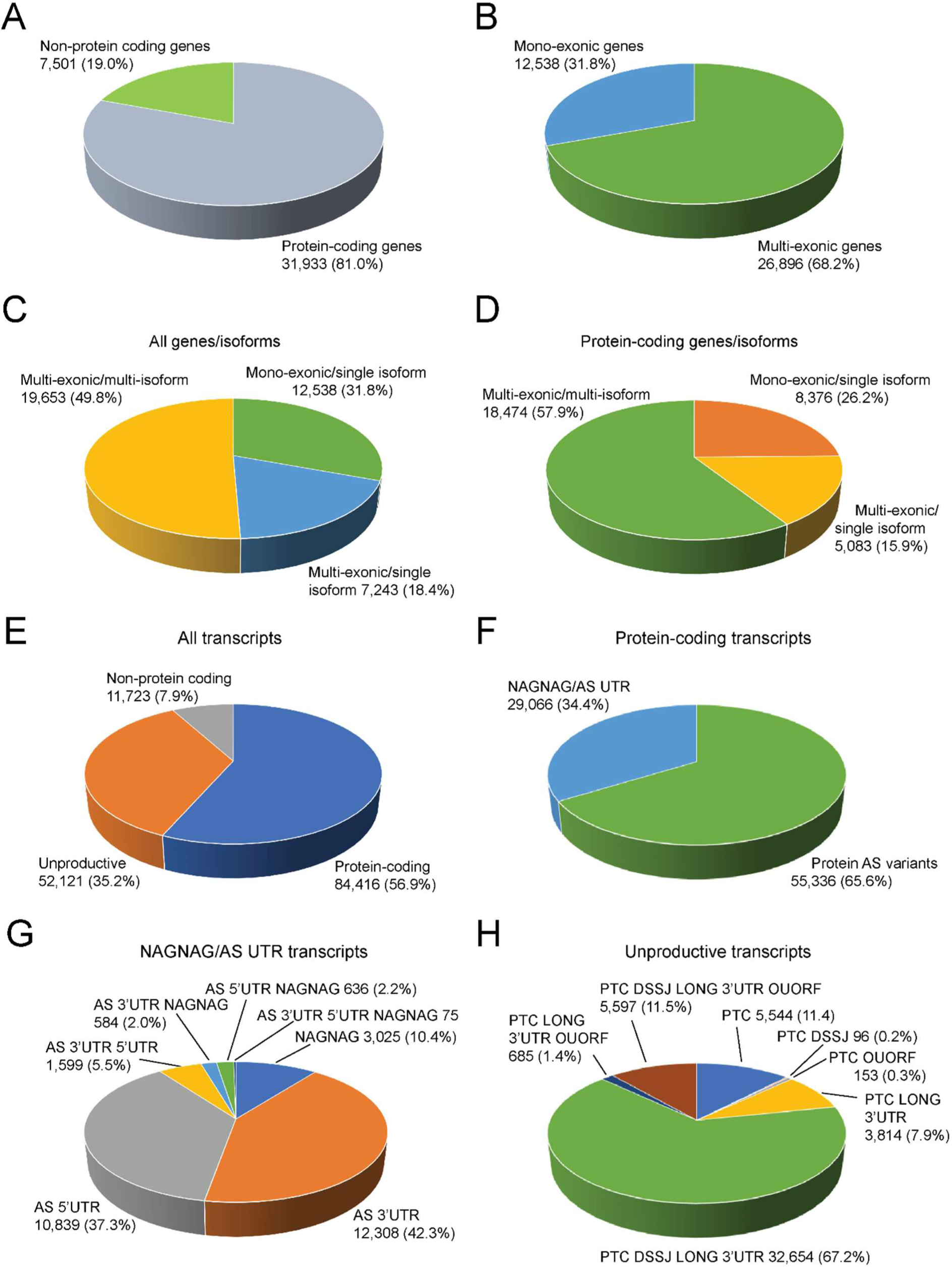
TranSuite analysis of BaRT2.0 transcriptome. The 39,434 genes of BaRT2.18 are divided into A) protein-coding and non-protein-coding genes; and B) mono-exonic and multi-exonic genes. The distribution of isoform numbers in mono-exonic and multi-exonic genes is provided for C) all genes, and D) protein-coding genes. The 148,760 transcripts of BaRT2.18 are divided into E) transcripts from protein-coding genes (both protein-coding and unproductive transcripts) and those from non-protein-coding genes; F) protein-coding transcripts are separated into those transcripts with no or little change in protein sequence due to AS occurring only in the 5’ and/or 3’UTR or NAGNAG AS which alters the coding sequence by a single amino acid and those coding for protein variants; G) NAGNAG and AS UTR transcripts are portioned in different combinations of AS events; H) unproductive transcripts which contain PTCs are divided by the presence of different NMD features (DSSJ – downstream splice junction; Long 3’UTR; OUORF – overlapping uORF). The analysis is based on gene classification using a minimum of 100 amino acids to define a protein-coding transcript.

At the transcript level, 136,537 (92.1%) BaRTv2.18 transcripts came from protein-coding genes. Of these, 61.8% encoded protein isoforms while 38.2% were unproductive (Figure 6E; Supplementary Table 11). Alternatively spliced transcripts that coded for proteins were divided into those where the AS events had little or no effect on the coding sequence (e.g NAGNAG AS sites or AS in the UTR) (34.4%) and those that encoded protein variants (65.6%) (Figure 6F; Supplementary Table 11). NAGNAG AS events generate transcripts that code for protein variants differing by only one amino acid and transcripts of genes where AS events occur only in the 5’ and/or 3’ UTRs code for identical proteins. Breaking these down further (Figure 6G; Supplementary Table 11) the most frequent AS events were in the 3’UTR (42.3%) then 5’UTR (37.3%) and NAGNAG events (10.4%), with lower numbers of combinations (Figure 6G). Across all transcripts, NAGNAG AS events were present in 2.9%. Finally, the unproductive transcripts from protein-coding genes were classified by their NMD features: presence of a premature stop codon (PTC), downstream SJs, long 3’UTR, or upstream ORF overlapping the authentic translation start site (67), (Figure 6H; Supplementary Table 11). 67.2% of the unproductive transcripts contained the classical combination of NMD features of a PTC with downstream splice junctions and long 3’UTRs with a further 20.9% having one or two of these features. Overall, 13.3% of unproductive transcripts contained an overlapping uORF (Figure 6H; Supplementary Table 11).

### BaRTv2.18 represents a barley transcript dataset with improved integrity

To evaluate BaRTv2.18, six benchmarks were compared between BaRTv2.18 and BaRTv1.0 (Supplementary Table 12). The highly conserved BUSCO gene set (59) was used to assess transcriptome completeness. BaRTv2.18 outperformed BaRTv1.0 in BUSCO benchmarking results, with 1,530 complete BUSCO genes compared to 1,501 in BaRTv1.0. BaRTv2.18 also contains fewer fragmented genes, with 38 compared to 70 in BaRTv1.0 (Supplementary Table 12). No transcripts overlapping Ns in the genome or duplicated sequences were identified in BaRTv2.18 (Supplementary Table 12) as they were removed during the filtering process (Figure 1). To assess the relative level of chimeric transcripts in BaRTv1.0 and BaRTv2.18, transcripts were compared to barley Haruna Nijo full-length cDNAs (flncDNAs) (61) and the percentage of transcripts that overlapped with multiple flcDNAs determined. BaRTv2.18 contained ca. 250 chimeric transcripts, derived from Iso-seq. This was lower (1.3%) than in BaRTv1.0 (2.23%) (Supplementary Table 12) reflecting more possible artefactual chimeric transcripts caused by short read transcript assembly. Thus, BaRTv2.18 outperforms BaRTv1.0 in the completeness and diversity of this new barley reference transcriptome.

### BaRTv2.18 represents an improved barley transcript dataset for alternative splicing analysis

The impact of BaRTv2.18 vs. BaRTv1.0 on transcript quantification accuracy was investigated by comparing splicing ratios of AS transcript isoforms using high resolution RT-PCR (HR RT-PCR) and those obtained directly from transcript quantification using RNA-seq data, different RTDs and Salmon. We used BaRTv1.0-QUASI (a ‘padded’ version of BaRTv1.0 (19)) which was constructed using the cv. Morex genome and BaRTv2.18 based on the cv. Barke as comparative RTDs. To evaluate whether BaRTv2.18 can be used to accurately quantify RNA-seq data beyond the Barke cultivar, HR RT-PCR data was generated using 85 (Morex derived) and 42 (Barke derived) primer pairs to amplify across known AS events and total RNA from both Morex (15 samples) or Barke (12 samples), respectively. Comparisons between RT-PCR data in Morex and Barke are compared with quantifications using Morex and Barke RNA-seq (Figure 7). The Morex HR RT-PCR dataset had more primer pairs, RT-PCR products and therefore more datapoints (Figure 7B and D). Using Barke RNA-seq, BaRTv2.18 showed superior performance over BaRTv1.0 producing higher Pearson and Spearman correlation coefficients between the splicing ratios derived from HR RT-PCR and RNA-seq (0.833 and 0.840, respectively) than BaRTv1.0-QUASI (0.777 and 0.780, respectively) (Figure 7A, C and E, Supplementary Table 13). Despite using Morex RNA-seq, BaRTv2.18 still achieved higher Pearson and Spearman correlation coefficients over BaRTv1.0, which is a Morex based assembly (Figure 7B, D and E, Supplementary Table 13). In summary, these results show the improvement of transcript models in BaRTv2.18 has led to a modest improvement in quantification accuracy when compared to BaRTv1.0, taking into account differences between cultivars Morex an Barke.

**Figure 7:**
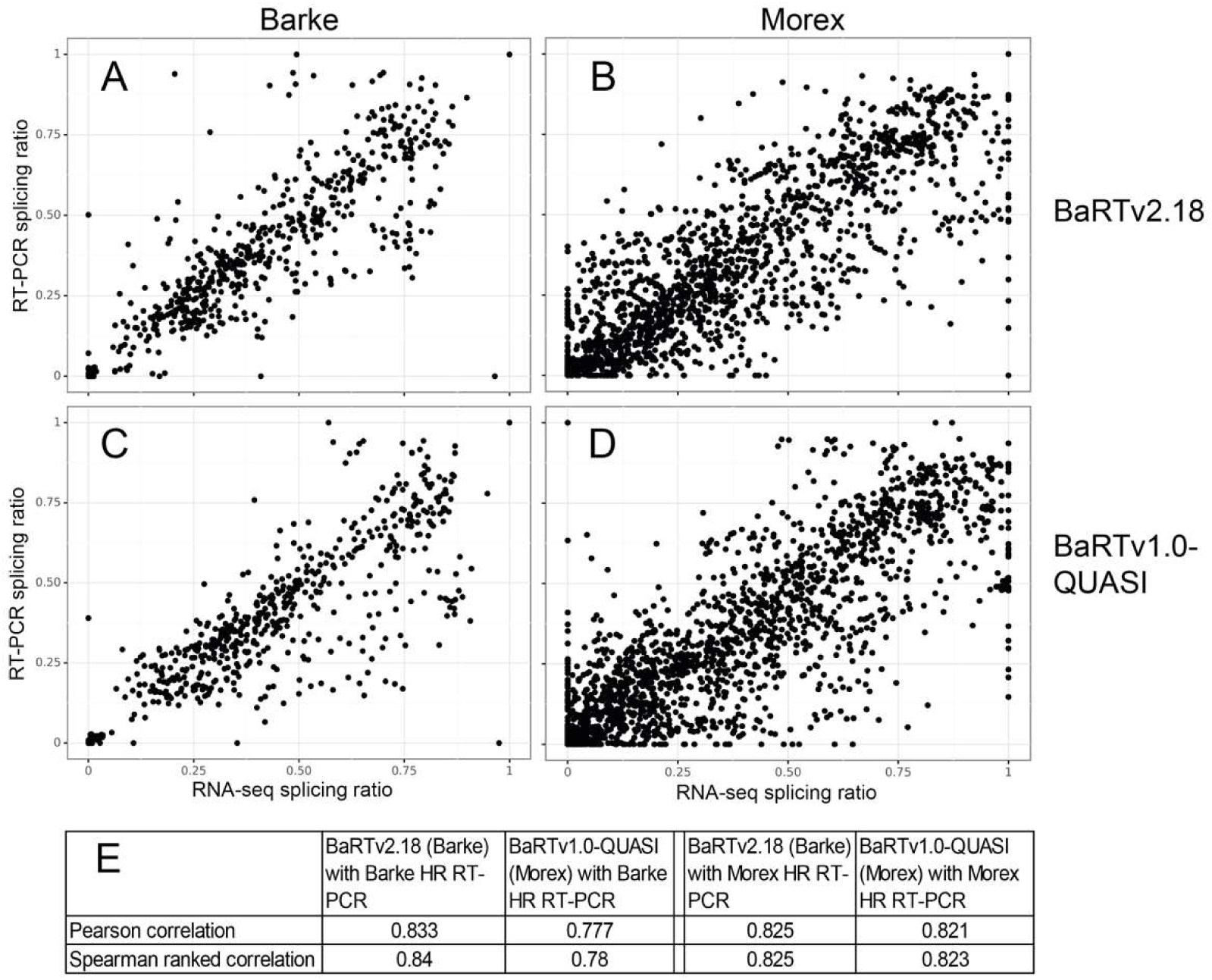
Comparison of accuracy of transcript quantification using BaRTv2.18 and BaRTv1.0-QUASI measured by correlation of splicing ratios from HR RT-PCR and RNA-seq. Splicing ratios were calculated for multiple AS events from different genes from relative fluorescence units from HR RT-PCR and transcript abundances (TPM) from RNA-seq data were quantified with Salmon. **A and B**) BaRTv2.18 was used to calculate splicing ratios from RNA-seq data from 12 samples of Barke and 15 samples of Morex RNA and compare these to splicing ratios from Hr RT-PCR using 42 and 85 primer pairs, respectively and the same RNA samples. **C and D**) The same was carried out using BaRTv1.0-QUASI. Pearson and spearman correlations for each of the combinations are shown (**E**).

### BaRTv2.18 represents a substantial improvement in defining transcript start and end sites

The predominant difference between BaRT2 and the short read based BaRTv1.0 (21) is the high quality Iso-seq forming an backbone with 94,247 (63.6%) transcripts from 20,969 (53.2%) genes (Supplementary Table 14). These were supported and complemented by stringently quality-controlled transcripts from the short-read Illumina assembly. An important feature of BaRTv2.18 that is provided by the Iso-seq dataset are accurately defined TSS and TES. Out of the 20,969 genes with Iso-seq transcripts with defined TSS/TES, we identified 47,750 and 62,468 TSS and TES respectively, an average of 2.28 TSS and 2.98 TES per gene. The majority of these genes have alternative TSS or TES; 52.5% (11,023 genes) have two or more TSS, whilst 63% (13, 217) of genes have two or more TES (Supplementary Figure 3).

The accuracy of the TSS and TES identified in the Iso-seq were validated in two ways. Firstly, TSS and TES were validated in silico by scanning the region around transcript 5’ and 3’ ends for enrichment of transcription motifs (TATA box, Initiator – Inr and Y patch), 3’-end processing signals (3’ polyadenylation signal – PAS and CFlm) and translation start sites (Kozak) (Supplementary Table 2; Figure 5). The positions of these motifs were compared using the transcript 5’ and 3’ ends from the BaRTv2.18 and BaRTv1.0 datasets. The TATA box is a T/A rich cis-regulatory element 25-35 bp upstream of transcription start sites that specifies where transcription begins (34, 68), while the Initiator (Inr) motif is pyrimidine-rich and overlaps the transcription start site and is important for activation of transcription (69). The Y patch is a pyrimidine rich sequence upstream of the transcription start site unique to plants and found in more than 50% of annotated rice genes (70), while the Kozak motif is found downstream of the transcription start site and includes the start AUG codon for translation initiation (71). The 3’ polyadenylation signal (PAS) motif is required for 3’ end polyadenylation (72), while the CFlm motif is the binding site of cleavage factor Im, an essential 3’ processing factor (73, 74). We found that the 5’ transcript ends from BaRTv2.18 were more enriched for TATA box, Inr, Y patch promotor motifs and the Kozak translation start site motif compared to BaRTv1.0 (Figure 5A-D). The TATA box motif presented a peak density of 0.055 instances per site (5,002 instances out of 91,123) ∼30bp upstream of TSS site in BaRTv2.18, compared to a maximum peak of 0.014 (1,844 instances out of 129,835) in BaRTv1.0 (Figure 5A), approximately a fourfold enrichment (proportion test p<2.2e-16), suggesting TSS positions in BaRTv2.18 are much more characteristic of a true TSS. The Inr and Y patch sequence motif showed similar results with 0.01029 and 0.0128 instances per site (938 and 1,170 instances out of 91,123) in BaRTv2.18 and 0.00642 and 0.00791 instances per site (834 and 1,027 instances out of 129,835) in BaRTv1 (proportion test p<2.2e-16). The Kozak motif had a defined peak at ∼150 nt downstream of the TSS at 0.0035 instances per TSS (322 instances out of 91,123) in BaRTv2.18 (Figure 5D) while BaRTv1.0 had a much broader distribution, with a peak of 0.0013 instances per TSS (168 instances out of 129,835). As a result, the Kozak motif is significantly enriched in BaRTv2.18 (proportion test p<2.2e-16). Similar results were obtained for PAS and CFlm 3’ end processing motifs with greater enrichment in BaRTv2.18 compared with BaRTv1.0, with a peak of 0.062 instances per TES (6,652 instances out of 107,977) for the CFlm motif in BaRTv2.18 compared with 0.038 (4,875 instances out of 129,928) in BaRTv1.0 and a peak of 0.019 instances per TES (2,095 instances out of 107,977) for the PAS motif compared with 0.0075 (969 instances out of 129,928) in BaRTv1.0 (Figure 5E, F). Proportion test for both TES related motifs shows significant enrichment in BaRTv2.18 (p<2.2e-16). These results strongly support the determination of TSS and TES by the methods used to analyse Iso-seq data and give confidence to the identification of HC TSS/TES.

Secondly, to confirm enriched TSS determined for Iso-seq transcripts, four genes with evidence of alternative TSS were selected for analysis by 5’ RACE. Sequencing determined the positions of 5’ RACE products for each of the four genes. These products matched Iso-seq TSS sequences exactly or were in close proximity (within 5bp) (Supplementary Figure 2). For example, BaRT2v18chr5HG238000 5’ RACE confirmed two of the three TSS including the most prominent TSS expressed in inflorescence and peduncle (Supplementary Figure 2A). Similarly, 5’ RACE products coincided with predicted TSS in BaRT2v18chr1HG022010 and BaRT2v18chr6HG297070 (Supplementary Figure 2B and C) and BaRT2v18chr6HG314960 (5’ RACE site 20 bp from most abundant TSS - not shown). Thus, 5’ RACE analysis supported predicted TSS, often with exact matches, for the genes examined.

### BaRTv2.18 allows the investigation of transcript isoform expression and differential TSS and TES usage across barley tissues and organs

To illustrate the utility of BaRTv2.18, RNA-seq data from the twenty different biological samples used in its construction were quantified using Salmon with BaRTv2.18 as the reference transcriptome. The full dataset of transcript quantifications is found in Supplementary Table 15. Analysis of BaRT2v18chr3HG130540 illustrates different levels of expression of AS isoforms and differential usage of alternative TSS and TES among different tissues/organs (Figure 8). BaRT2v18chr3HG130540 is a 59 kD U11/U12 small nuclear ribonucleoprotein component of the spliceosome and is alternatively spliced to give nine Iso-seq transcript isoforms. The gene has two TSS and four TES (Figure 8A). Five of the transcripts appear to code for protein (.1, .2, .6, .7, .8) (Figure 8A) while four are unproductive in containing PTCs and NMD features (.3, .4, .5 and .9) (not shown). The composition and relative abundances of different transcript isoforms clearly differ among the various samples (Figure 8B), and the expression of some transcripts is limited to specific tissues. For example, BaRT2v18chr3HG130540.6 has an intron in the 3’UTR and appears to be the predominant isoform expressed in six-day-old embryonic tissue, heat-stressed coleoptiles and peduncle with low or no expression in other tissues (Figure 8B; Supplementary Table 15). The other protein-coding transcripts .1, .2, .7 and .8 code for the same protein but have alternative TSS and TES (Figure 8A). The .1 and .2 transcripts have the same TSS but different TES (Figure 8A) while the .7 and .8 pair have the same TSS (different from the TSS in .1 and .2) and different polyadenylation sites. Most samples expressed predominantly one or other of these pairs of transcripts while tissues which expressed both usually expressed one more highly (Figure 8B). For example, while .7 was expressed in most tissues, the .8 transcript was found mainly in coleoptiles, day 55 lodicules, day 55 palea and 6-day-old embryos (Figure 8B). Variation in the levels of, for example, .2 and .8 in different tissues illustrates differential usage of alternative TSS. Similarly, variation in abundance of .1 and .2 transcripts among different tissues reflect differential usage of alternative TES. The PTC-containing unproductive transcripts were generally less abundant than the other transcripts reflecting their likely sensitivity to NMD. These transcripts had either skipping of exon 6 or intron retention events. Thus, using BaRTv2.18 different transcript isoforms are readily detected and can be quantified at both the gene and transcript level.

**Figure 8:**
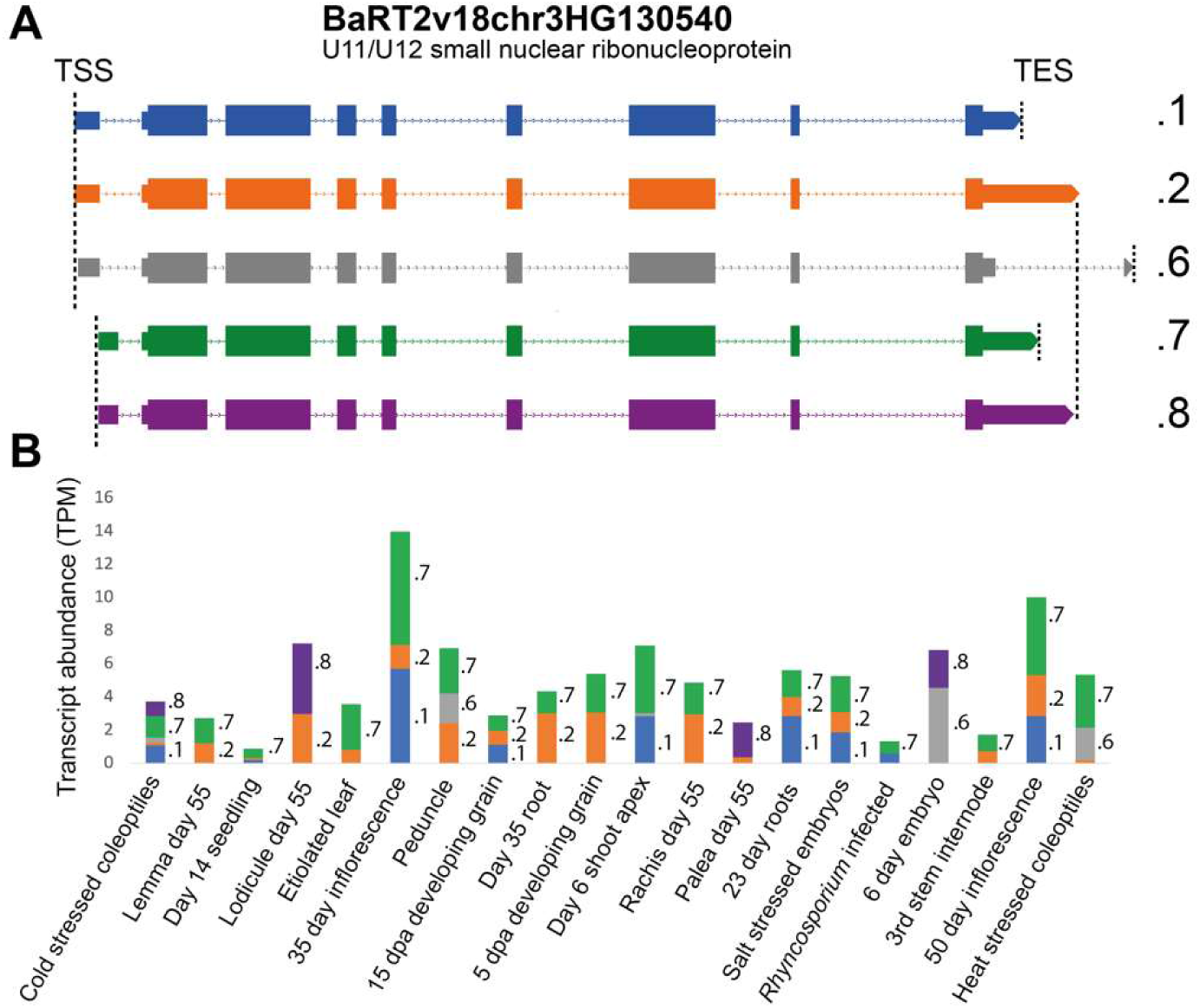
Differential TSS and TES usage in a range of barley tissues and conditions. Transcript abundance of transcript isoforms from gene BaRT2v18chr3HG130540, annotated as a U11/U12 small nuclear ribonucleoprotein 59 kDa protein across the 20 samples used to construct BaRTv2.18. **A**) Transcript structures of protein-coding isoforms; dotted lines - introns, thick lines - UTR exons and very thick lines - CDS regions. **B**) Relative abundance of different isoforms (TPM) for each of the 20 samples are shown, with individual transcript quantifications indicated by different colours and labelled with isoform number within each bar.

## DISCUSSION

The potential of RNA-seq experiments is often not fully realised due to the widespread lack of high-quality transcriptome annotations (6, 20, 21). Stringently filtered, comprehensive RTDs permit fast and accurate gene and transcript level quantifications to be obtained from ultra-deep short read RNA-seq data (2, 11, 20) using Salmon and Kallisto (15, 16). Here we present a new barley RTD resource, BaRTv2.18, based on parallel Iso-seq and Illumina data collected exclusively from the 2-row spring barley cv. Barke, a genotype directly relevant to present-day European agriculture and contemporary germplasm (11, 21, 41). The recently released chromosome-scale TRITEX assembly of the Barke genome (Figure 2A, (41)) was used as the reference for mapping the Barke RNA-seq data. The previously published barley RTD, BaRTv1.0, was constructed from a diverse collection of over 800 short read RNA-seq samples in 11 datasets including different developmental stages and abiotic stresses from a range of barley cultivars and landraces using the genome sequence of barley cv. Morex as reference (21). Although BaRTv1.0 may contain greater transcript diversity (total of 177,240 transcripts) due the range of genetic lines used, 5’ and 3’ ends of transcripts BaRTv2.18 therefore provides significant improvements over BaRTv1.0: 1) The majority of BaRTv2.18 reference transcripts (94,247-64%) are derived from Iso-seq reads and have accurate TSS and TES as well as SJs. 2) the defined TSS and TES allow investigation of alternative TSS and TES usage reflecting enhanced resolution of transcriptional and post-transcriptional gene regulation not possible with BaRTv1.0; 3) accurate transcript quantification using BaRTv2.18 does not require padding of transcripts (artificial extension of different transcripts of a gene to the length of the longest transcript - needed for BaRTv1.0 as 5’ and 3’ ends were not defined – Ref 21); and 4) SJs, TSS and TES can now be determined single molecule sequencing data without the need for parallel experimental end determination techniques or hybrid correction. Iso-seq has been widely used to address transcriptome diversity in plant and crop species. However, issues still exist with false SJs caused by mapping errors around the introns, incomplete gene and transcript coverage and accuracy of determination of TSS and TES. Methods to correct sequence errors impact transcript accuracy as both self-correction and hybrid correction of errors can lead to loss of AS events and introduce novel errors (over-correction). We have used the new Iso-seq analysis pipeline based on that carried out for Arabidopsis AtRTD3 (43) with some modifications for the barley dataset used. We used TAMA for greater control of analysis, a splice junction centric approach to identify and remove false SJs, and a probabilistic approach to determine TSS and TES which take transcript abundance into account. Although the latest PacBio HiFi reads contain lower error rates (75), false positive novel isoforms will still be an issue for CCS reads with a low number of passes. These problems are addressed by our pipeline. Short read transcript assembly is used to provide gene and transcript coverage for gene regions where Iso-seq has little or no coverage. We applied a new software pipeline (RTDmaker) for stringent quality control of transcripts; RTDmaker filters transcripts to remove, for example, redundant and fragmentary transcripts and retain only transcripts with high quality SJs. The combination of these methods ensured that BaRTv2.18 represents the most accurate and diverse barley transcriptome to date.

Other pipelines, besides the PacBio Iso-seq analysis pipelines, have been employed to improve determination of transcript features. For example, the Integrative Gene Isoform Assembler (IGIA) for cotton (8) combined both Iso-seq and RNA-seq to provide maximum coverage of genes and transcripts. The study also used CAGE-Seq and PolyA-Seq to define TSS and TES, respectively which were used to correct Iso-seq sequence 5’ and 3’ ends (8). The number of CAGE-seq and PolyA-seq sites over the expression threshold were found in 22,863 and 23,736 genes, respectively, representing around one third of cotton genes (8). Here, 20,969 barley genes have Iso-seq support, each with both TSS and TES defined. Importantly, our method does not require experimental determination of ends by CAGE-Seq or PolyA-Seq data, rather it maximises the information that can be obtained from Iso-seq by determining TSS and TES directly from the Iso-seq dataset using our novel approaches. Furthermore, our method does not require hybrid error correction and so avoids over-correction (51); instead, Iso-seq transcripts are removed if they are not supported by high confidence SJs, TSS and TES. The accuracy of TSS and TES for the Iso-seq derived transcripts in BaRTv2.18 was shown by the increased enrichment of characteristic sequence motifs of known TSS, TES and translation start site sequence motifs in gene sequences in BaRTv2.18 in comparison to BaRTv1.0 (Figure 5) and by 5’RACE of specific genes. In plants, TSS have been identified in Arabidopsis, maize and cotton (8, 34, 35) and TES in Arabidopsis and cotton (8, 39, 40). CAGE-Seq analysis in maize has shown that ∼70% of annotated gene models have mis-annotated TSS, reflecting the inaccuracy of short read annotation (35). In many cases, the CAGE TSS mapped downstream of annotated TSS suggesting that the short read assembled transcripts were artificially extended as seen for Arabidopsis in TAIR10 and Araport (20). Comparing enrichment of promotor motifs between BaRTv1.0 and BaRTv2.18 confirmed the expected enhanced definition of TSS and TES in BaRTv2.18 (Figure 5). Quantitative CAGE-Seq in cotton found that 40% of gene loci had alternative TSS (8) whilst, in maize, hundreds of significant differences in promotor usage were identified between tissues (35). Here, in the genes with high confidence TSS and TES sites, over a half had two or more TSS or TES sites. The advantage of BaRTv2.18 having well-defined TSS and TES for many genes is that differential TSS and TES usage can be addressed in RNA-seq analysis, as illustrated by BaRT2v18chr3HG130540 showing tissue-specific variation in alternative TSS or TES usage (Figure 8).

More than half of the transcripts in BaRTv2.18 are based on HC Iso-seq and the genes with low or no Iso-seq coverage required complementation with short read assembled transcripts. These transcripts have been generated by multiple assemblers to capture transcript diversity and stringent quality control to remove false transcripts and retain only well-supported transcripts. The accuracy of assembly is illustrated by the high number of short read transcripts which are replaced by Iso-seq transcripts during merging of BaRTv2.0-Iso and BaRTv2.0-Illumina. Therefore, although the 5’ and 3’ ends of these transcripts should be treated with caution, their SJs are accurate and are a valuable contribution to BaRTv2. As further improvements in single molecule sequencing technology are made to significantly improve Iso-seq depth of coverage, ultimately, complementation with short read assemblies will not be required. Similarly, our methods avoid the need for hybrid error correction which also helps single molecule sequencing analysis to move towards being a self-contained process where complete transcript characterisation is achieved directly from the data.

BaRTv2.18 improved transcript quantification from RNA-seq data as demonstrated by the high correlations of values of AS splicing ratios compared to BaRTv1.0 (Figure 7; Supplementary Table 13). In this comparison, the QUASI (padded) version of BaRTv1.0 was compared to the unadjusted BaRT2.18 version. We showed previously in Arabidopsis AtRTD2 that variation in the 5’ and 3’ ends of transcripts from the same genes could skew transcript quantification and that padding of transcripts to the length of the longest transcript overcame this problem (20); the BaRTv1.0-QUASI version similarly improved transcript quantification in barley (21). The accurate determination of TSS and TES for many transcripts and the removal of short, fragmented transcripts in BaRTv2.18 should overcome the need for padding of the RTD. The unpadded BaRTv2.18 generated more accurate quantification than BaRTv1.0-QUASI as shown by the correlation of splicing ratios from HR RT-PCR and RNA-seq. There was also an effect of cultivar source on the correlations where comparing splicing ratios from HR RT-PCR and RNA-seq from the same cultivar had higher correlation values. This suggests that higher quantification accuracy is achieved when using the same genotype in RNA-seq that has been used to generate the RTD and will influence the development of a new pan-transcriptome to complement the recently published pan-genome (41). Comparative benchmarking also confirmed that BaRTv2.18 provides a substantial improvement over BaRTv1.0 (Supplementary Table 12). The reduction in fragmented BUSCO genes in comparison to BaRTv1.0 most likely reflects both the improved transcriptome per se and the higher quality of the Barke genome in comparison to the Morex genome that was used with BaRTv1.0 for read mapping (41, 62). In other metrics, BaRTv2.0-Iso performed well, with relatively few duplicated sequences (35) despite no specific efforts to remove them, and low numbers of possible chimeras (0.3%).

High quality RTDs form an important community resource for understanding the regulation of gene expression. Therefore, the BaRTv2.18 RTD is also supported by extensive transcript characterisation and annotation (Suppl. Tables 16, 17 and 18). Accurate translations of each transcript have been annotated using Transuite (52). The program fixes the authentic translation start site of each gene and translates all the transcripts from the fixed AUG. This overcomes mis-annotation of ORFs and distinguishes protein-coding and unproductive isoforms (20, 76). Transcripts are also annotated with potential functions and co-ordinates of transcript features such as CDS, exons, 5’ and 3’ UTRs are provided. The information should enable accurate interpretation of expression changes and allow direct comparisons among transcriptomics experiments and integration of data analyses conducted in diverse organisations. The methods that we have developed for both Iso-seq and Illumina analysis will allow any new long and short read datasets to be efficiently added as they become available and will improve the construction of RTDs from other cultivars and feed into development of a barley pan-transcriptome. Clearly, to maintain their community value, improvements to transcriptome resources should evolve alongside parallel improvements to emerging genomic resources (41, 62, 77, 78). Importantly, BaRTv2.18 will play a valuable role in providing experimental support for annotation of the most recent barley genome assemblies.

In summary, BaRTv2.18 represents a new and improved Reference Transcript Dataset built upon a broad framework of highly informative Pacbio Iso-seq full length transcripts incorporating accurate SJ, TSS and TES information. It is based upon the European 2-row spring barley cv. Barke which itself is highly relevant to contemporary barley agriculture and current research. The combination of the novel Iso-seq analysis and filtering pipeline, the new automated quality control pipeline for short read assembled transcripts and detailed transcript characterisation and annotation make BaRTv2.18 a high-quality community-wide reference dataset.

## AVAILABILITY

The BaRTv2.18 transcriptome and annotation files are available at https://ics.hutton.ac.uk/barleyrtd/bart_v2_18.html. Scripts used to produce BaRTv2.18 are available at the repository https://github.com/maxecoulter/BaRT-2. Raw Illumina and Iso-seq datasets have been uploaded to the Sequence Read Archive under the accession numbers <u>PRJNA755156</u> and <u>PRJNA755523</u>.

## SUPPLEMENTARY DATA

## ACKNOWLEDGEMENT

We thank Susanne König and Axel Himmelbach from IPK Gatersleben for preparing and sequencing the Iso-seq libraries

## AUTHOR CONTRIBUTIONS

RZ, JWSB, RWa, NS and GM designed experiments. MC, JCE, WG, RWo, NM, CS carried out experiments. MC, JCE, MB, MS, CS, JWSB, WG and RZ analysed data. LM maintained and updated the BaRT annotation website. The manuscript was written by MC, RZ, JWSB and RWa with contributions from all other authors.

---

**Supplementary Figure 1:**
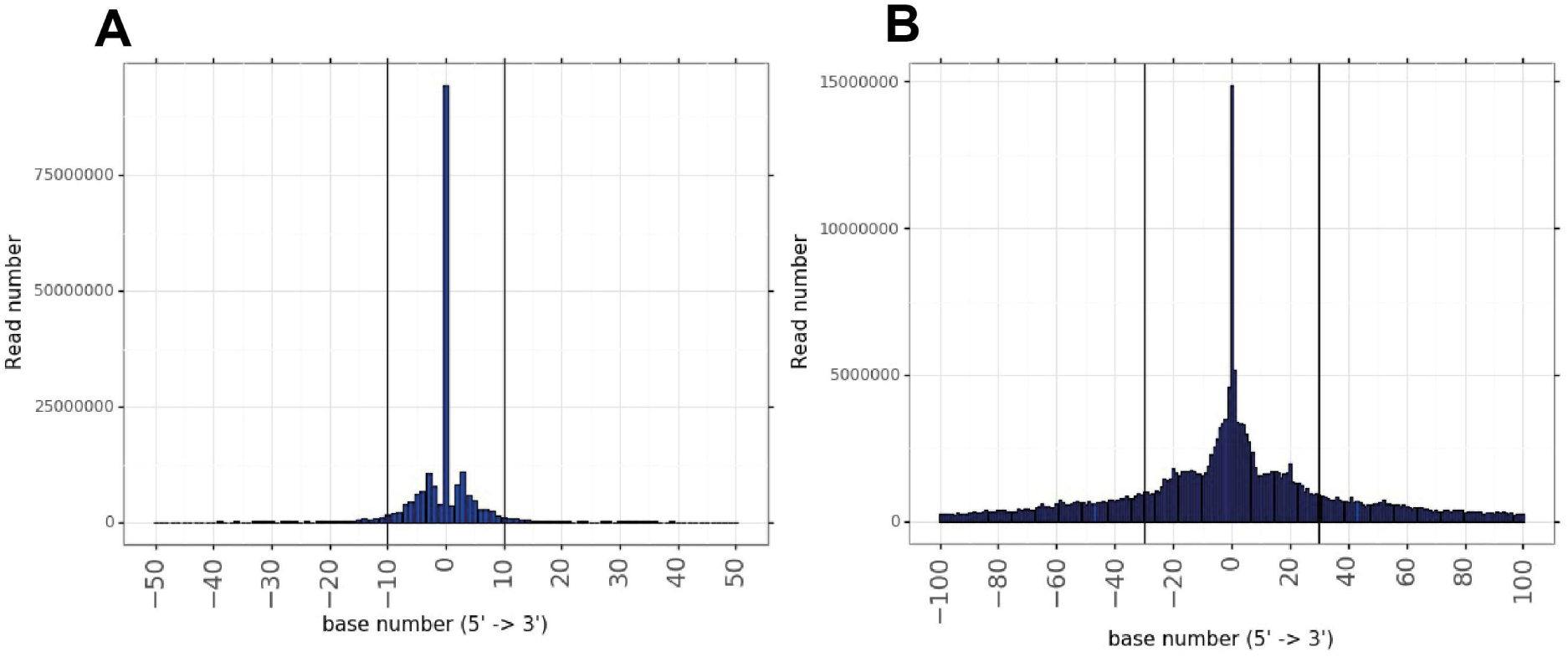
Histogram of end read support to determine window sizes for filtering of Iso-seq dataset. **A** distribution of 5’ read ends around 5’ ends of transcripts with high coverage (>= 10 reads), **B** distribution of 3’ read ends around 3’ ends of transcripts with high coverage (>= 10 reads). Window sizes used for filtering 5’ and 3’ ends of transcripts are indicated in C (+/-10) and D (+/-30) by vertical lines.

**Supplementary Figure 2.**
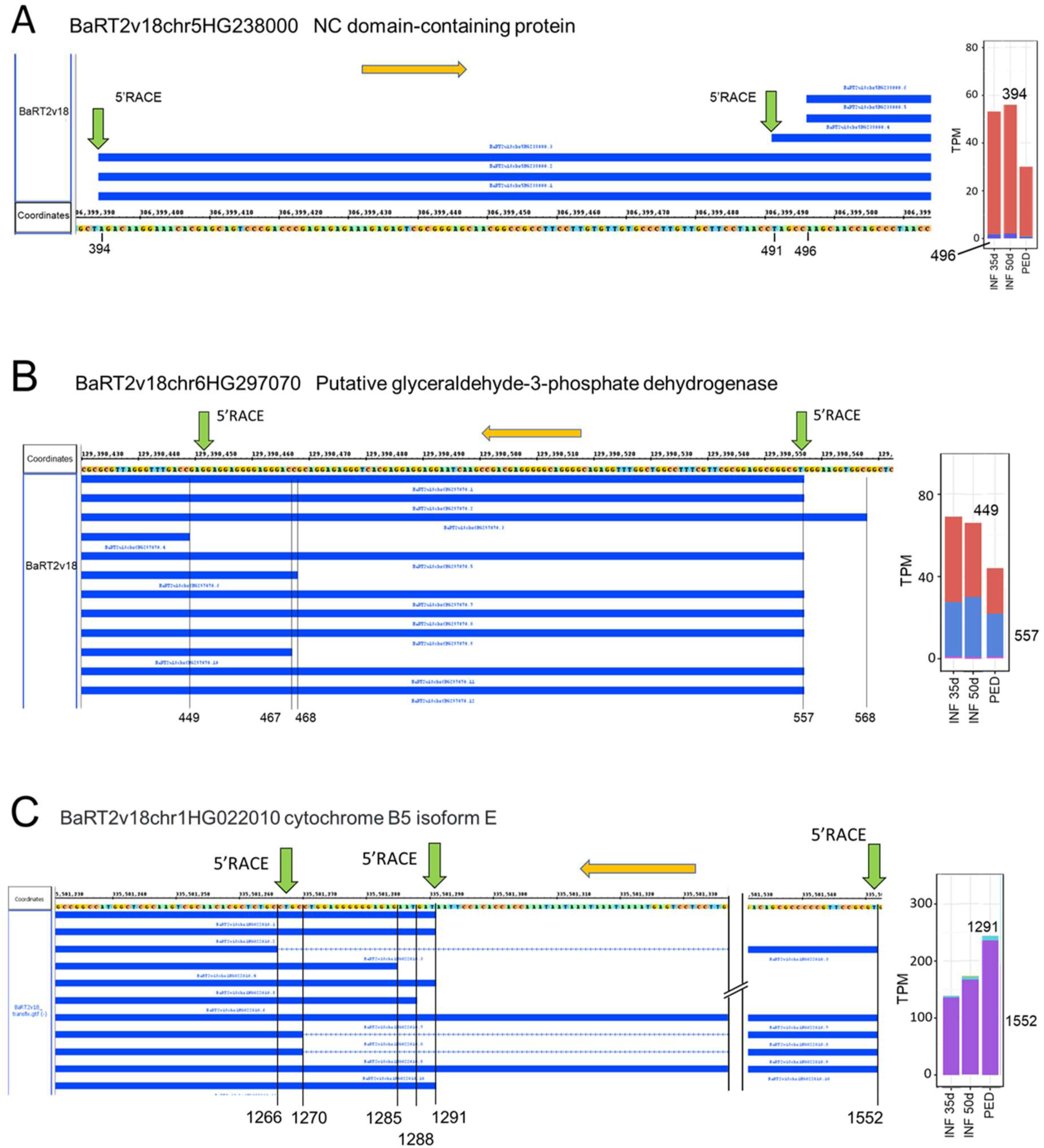
Confirmation of predicted TSS by 5’RACE.5’ RACE. **A**) BaRT2v18chr5HG238000. TSS of transcript isoforms are visualised on Integrated Genome Browser (IGB) along with the genomic sequence and the last digits of the co-ordinates are indicated. 5’ RACE products are shown by green arrows. The right hand panel shows the relative abundance (TPM) of expression of isoforms from the different different TSS in inflorescence (INF) and peduncle (PED) tissues. Orange arrow – direction of transcription. **B**) BaRT2v18chr6HG297070 – the two 5’ RACE sites correspond to the most abundant transcripts (legend as in A). **C**) BaRT2v18chr1HG022010 has a TSS (1552) 250 bp upstream of a second TSS (legend as in A). The 5’ RACE signal at 1552 coincides exactly with the major TSS. No signal was detected at 1291 but transcripts using this TSS are much more abundant in other tissues (e.g. roots, rachis, internode – not shown).

**Supplementary Figure 3:**
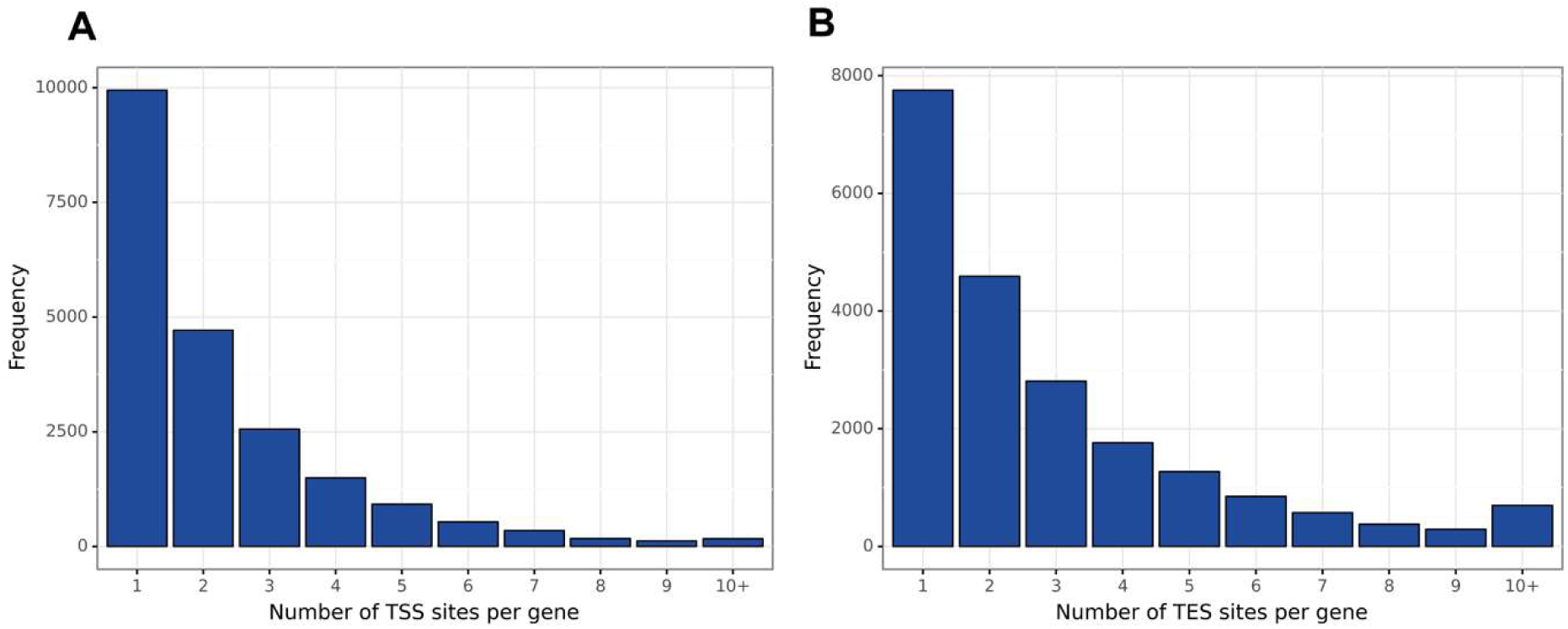
Histogram of TSS (A) and TES (B) per gene in BaRTv2.18 genes with Iso-seq support

