## Supplementary material for "BaRTv2: A highly resolved barley reference transcriptome for accurate transcript-specific RNA-seq quantification": supp materials and methods

**Supplementary materials and methods**

**Plant material**

*Seed samples:* All seed was sterilised by soaking in 70% ethanol for five minutes and then in a solution of 10% bleach with 0.1% Tween for ten minutes. Sterile distilled water (SDW) was added, and seeds were left for five minutes, followed by four more 1 min SDW washes.

*Vegetative tissues:* Leaf tissue was sampled at 14 days after planting (dap), while the etiolated leaf sample was grown in the dark and taken at 11 dap. Root samples were taken at 23 and 35 dap. Whole embryo (including developing root and coleoptile) and shoot apex samples were obtained from seeds grown on petri-dishes and sampled at 6 dap (the shoot apex was defined as the white base of the shoot). The shoot apex sample was from 5 developing embryos and the 3^rd^ stem internode tissue was dissected at 44 dap.

*Developing inflorescence samples:* samples were taken at 35 dap and 50 dap. At 55 dap, inflorescences were dissected to isolate lemma, palea, rachis and lodicule samples. Developing grain was taken from inflorescences 5 and 15 days post anthesis; two central caryopses were taken from each plant, making six developing grains per sample. Peduncles were taken when they had reached 3 – 5 cm in length.

*Heat and cold stress:* coleoptiles were dissected from seed grown in dark on petri-dishes. Three petri-dishes were used for each cold stress and hot stress sample. For cold and heat stress, petri-dishes were kept in a fridge and incubator at 4°C and 30°C respectively for 48 hours. Three coleoptiles were taken at 0h (before temperature change), 2 h, 4 h, 24 h and 48 h post temperature change. For salt stress 4.35g of salt was added to 500mls distilled water agar (DWA) to make up a 150mM salt agar. Seed grown for seven days on plates of DWA was added to three petri-dishes of the salt agar. Three whole embryos (including developing roots and coleoptile) were taken at 0h (before salt stress), 4 h, 8 h, 24 h and 48 h.

*Biotic stress:* The *Rhynchosporium commune* isolate 214 gfp was grown on CZV8CM medium (40) for three weeks at 17°C in the dark. Fungal spores were harvested by scraping the mycelial mat with a spatula following the addition of 5 ml SDW. The mixture was added to a sterile filter column with 60 µm filter paper to filter spores. The resulting spore suspension was added to a falcon tube and centrifuged for 3 min at 1600 x g, washed twice with 5 ml of SDW, followed by centrifugation at 1600 x g for 3 min. The spore suspension was adjusted to a final concentration of 3 x 10^5^ spores/ml. 4.6 ml of spore suspension was sprayed on 22-day old seedlings. 10 µl drops were added to one leaf to observe growth of mycelia by confocal microscopy. Plants were kept in the dark for 24 h and sealed at 100% humidity for 48 h. Confocal microscopy showed penetration of hyphae at 3 dpi, so leaf samples were taken from 3 plants at 3 dpi and 9 dpi to reflect early and middle stages of infection (41).

Seven further tissues were collected at IPK Gatersleben. Plant material was grown in the greenhouse at IPK Gatersleben with day/night temperatures of 21/18^o^C. Embryonic tissue, leaves, roots, internode, inflorescence (5 mm), developing seeds (5 and 15 days after pollination) were collected as described before (42), snap frozen in liquid nitrogen and stored at -80^o^C until RNA extractions were performed.

**Iso-seq processing**

The 20 Iso-seq libraries were processed individually. Raw subreads were initially processed using the CCS tool from IsoSeq3 to create circular consensus sequences (CCS) with the parameters “ccs --min-rq 0.9 -j 28”, with –min-rq referring to the minimum accuracy for a read to be omitted, and -j flag the number of threads. The resulting CCS reads were stripped of adapter sequences and poly-A tails using lima and refine from Isoseq3, with parameters “--isoseq --peek-guess” for lima and default parameters for refine. A further clean-up of PolyA tails was carried out using the Transcriptome Annotation by Modular Algorithms (TAMA) tama_flnc_polya_cleanup.py script (<https://github.com/GenomeRIK/tama/wiki/TAMA-GO:-Sequence-Cleanup>).

The resulting full length non-chimaeric (FLNC) reads were mapped to the Barke genome (39) using Minimap2 (43) with the parameters “-ax splice:hq -uf -G 6000 --secondary=no”, with the “splice:hq” option recommended for mapping PacBio ccs reads, the “-uf” flag for prioritising canonical splice sites by transcript strand, and “-G” referring to maximum gap in nucleotides on the reference genome. Unmapped reads were removed using samtools with the parameters “view -h -F 4” (44).

The mapped FLNC reads were then processed into a .bed annotation file of genes and transcripts using TAMA collapse (<https://github.com/GenomeRIK/tama/wiki/Tama-Collapse>) with parameters “-d merge_dup -x capped -m 0 -a 0 -z 0 -sj sj_priority -lde 30 -sjt 30”, with “merge_dup” for merging duplicate transcripts, “-x” flag for whether 5’ capping was used, “-m”,”-a”,”-z” referring to SJ, start and end differences allowed between merged transcripts respectively. The “sj_priority” option is used for generating information on errors either side of SJs, while -Ide and -sjt flags refer to the number of errors around SJs allowed and size of window for observing these errors, meaning that only reads with large insertions or deletions close to SJs were removed. The local error density file produced using these parameters was used later for splice junction filtering. 32,260 reads in total failed TAMA collapse local_density_error threshold ("Ide_fail") and were removed. This left 6,211,392 FLNC reads that were used for gene and transcript annotation.

The 20 .bed files from TAMA collapse were merged together using TAMA merge (<https://github.com/GenomeRIK/tama/wiki/Tama-Merge>) with the parameters “-m 0 -a 0 -z 0 -d merge_dup”, with -m, -a and -z settings set to 0 in order to keep transcripts at single nucleotide resolution at this stage and only merge transcripts with same start, end and splice junctions. A small number of transcripts had exons which overlapped a gap in the genome (denoted by Ns). Transcripts containing Ns were removed as the Ns would cause mis-mapping. A .bed file containing the co-ordinates of gaps in the Barke genome was generated by custom code and transcripts overlapping these coordinates in the exons were identified and removed from the TAMA merge .bed file using bedtools intersect (45) with parameters “-split -v”. The original, unfiltered .bed file was used as well as the .bed file with transcripts with Ns removed as information on number of reads in each gene was required for downstream filtering.

**RTDmaker processing of Illumina transcriptome**

The RTD was quality controlled using RTDMaker (<https://github.com/anonconda/RTDmaker>). RTDmaker identifies and removes 1) redundant transcripts; 2) transcripts with splice junctions with low read support (3 reads in samples); 3) transcript fragments; 4) poorly supported antisense transcripts; 5) unstranded models; 6) antisense transcript fragments and 7) low expressed transcripts (cut-off default = 1 TPM). Redundant transcripts share exact intron co-ordinates and the longest transcript is retained. Redundant mono-exonic models are identified according to their genomic overlap. RTDmaker merges the remaining high-quality assembled models by identifying and removing redundant transcripts, transcript fragments and chimeric models. Transcript fragments were defined as being <70% of the longest transcript. RTDmaker then groups the transcript models according to their genomic co-ordinates and re-annotates their gene IDs. RTDmaker also generates a report detailing the characteristics of the resulting RTD, such as the number of genes, transcripts and isoforms. RTDmaker can also identify and remove transcripts from unstranded models and antisense transcript fragments. Antisense transcript fragments had a length that was <50% of the transcript on the opposite strand. Finally, RTDmaker performs a Splice-Junction (SJ) support analysis and a transcript abundance analysis. For the SJ-analysis, RTDmaker counts the number of unique reads that map to each SJ in each sample of the RNA-seq dataset (using the SJ log files generated by STAR). If a transcript contains a SJ that is not supported by the minimum required number of uniquely mapped reads in a minimum number of samples, then the transcript model is removed. For BaRT2.0-Illumina, a minimum SJ read support of 5 uniquely mapping reads in at least 1 sample (as there is only one sample per tissue) was required. Also, RTDmaker identifies low-expressed transcripts by quantifying transcript expression levels in the RNA-seq dataset using Salmon and transcripts with lower abundance than the minimum required (usually <1 TPM) in a minimum number of samples are removed. For BaRT2.0-Illumina, a minimum abundance of 1 TPM in at least 1 sample was applied. The number of rejected transcript models at each step of RTDmaker analysis is reported in Supplementary Table 8. Finally, a custom script (<https://github.com/maxecoulter/BaRT-2>) was used to remove 489 strandless transcripts and transcripts that overlapped Ns in the genome.

**RT-PCR analysis**

The RNA-seq data from the 12 Barke samples used for RT-PCR quantified using Salmon version 1.3.0. BLAST (blastn-short command, version 2.5.0+, (51)) was used to map the primers against the transcriptome. Primers that mapped to one location with perfect hits were used to identify transcriptome products that matched the RT-PCR products. As multiple transcripts can have the same AS event, the same size transcriptome products from each primer pair were clustered together. Only products >90bp and <700bp were used in the analysis. The clustered products were then matched to the corresponding RT-PCR products, with a window of ±6bp used to allow for errors in HR RT-PCR size calling. This matching was carried out computationally and checked manually. Primers that matched two or more transcriptome products that could be matched to HR RT-PCR products were used to estimate proportions of splice variants. Proportions of transcriptome products extracted from the RNA-seq data were estimated for each sample by taking the total TPM (transcripts per million) for each transcriptome product per primer set and dividing this by the sum of the TPM for all transcriptome products for that primer set. The same principle was applied to each HR RT-PCR primer pair, except here relative fluorescence unit (RFU) values for each product were divided by the total RFU to give proportions. Pearson and Spearman coefficients of correlation were calculated to determine how closely the AS proportions identified with RNA-seq and different transcript datasets matched the HR RT-PCR proportions. To compare the quantification accuracies between BaRTv1.0 and BaRTv2.0 the above method was used with BaRTv1.0-QUASI, and BaRTv2.0 with both 12 Barke HR RT-PCR/RNA-seq dataset and the Morex HR RT-PCR/RNA-seq datasets. To compare different components of BaRTv2.0, BaRTv2.0-Iso and BaRTv2.0-Illumina were also used for generating quantifications for correlating the 12 Barke RNA-seq/HR RT-PCR dataset.
